# Engineered Promoter System Enables High-efficiency Transgenic CRISPR Editing in Malaria Transmitting Mosquito *Anopheles sinensis*

**DOI:** 10.1101/2025.09.29.679135

**Authors:** Jun-Feng Hong, Qi-Li Zou, Xin-Yuan Xie, Yan-Ping Jiang, Si-Yi Wang, Xia Ling, Cao Zhou, Wei Sun, Xi Cai, Yong-Xin Yang, Yu Chen, Bin Chen, Liang Qiao

**Affiliations:** Chongqing Key Laboratory of Vector Control and Utilization, Institute of Entomology and Molecular Biology, College of Life Sciences, Chongqing Normal University, Chongqing, China; Laboratory of Evolutionary and Functional Genomics, School of Life Sciences, Chongqing University, Chongqing, China

**Keywords:** Promoter engineering, Transgenic system, CRISPR/Cas9 genome editing, Genetic modification, *Anopheles* mosquitoes

## Abstract

The CRISPR/Cas9 system deployed through crosses of transgenic lines expressing *Cas9* and *gRNA* facilitates efficient mutagenesis. However, its application in non-model insects remains limited, primarily due to a lack of well-characterized promoters capable of driving robust and stable expression of *Cas9* and *gRNA*. In the malaria mosquito *Anopheles sinensis*, we evaluated several ovary-biased promoters—*Asvasa2, Aszpg*, and *Asnanos*—for driving *Cas9* expression. Notably, the *Asvasa2* promoter mediated mutagenesis in nearly 60% of G_0_ individuals following microinjection of *gRNA*^*Aswhite*^. Among four RNA polymerase III promoters derived from *AsU6* genes, *AsU6*-1 yielded the highest *gRNA* transcriptional output, enabling 62% editing efficiency in G_0_ offspring. In addition, hybrid crosses between established transgenic lines demonstrated that the *Asvasa2-Cas9* and *AsU6*-1-*gRNA* combination enabled complete germline editing penetrance, where all F_2_ progeny inherited the intended mutations. This work provides a essential genetic toolkit for synthetic biology applications in *Anopheles* mosquitoes and a scalable framework for engineering other non-model insects.

---

Mosquitoes are major vectors of severe infectious diseases, including malaria and dengue fever, representing a persistent public health challenge worldwide[1]. Beyond their medical significance, mosquitoes serve as important systems for fundamental biological research, contributing to studies on chemosensation, swarming mating dynamics, host–pathogen interactions, and innovative genetic control strategies[2-4]. The CRISPR/Cas9 genome editing system has emerged as a pivotal tool for dissecting molecular mechanisms, evolutionary adaptations, and for developing novel genetic interventions in mosquitoes[5]. However, the conventional approach of microinjecting Cas9-*gRNA* ribonucleoprotein (RNP) complexes into embryos suffers from the absence of selectable markers, making the screening of edits inefficient in the absence of visible phenotypes. Moreover, this method is hampered by labor-intensive procedures and high embryonic mortality[6, 7]. Additionally, RNP-based editing lacks the spatiotemporal precision necessary for stage- or tissue-specific genetic modification.

In established model mosquitoes such as *Anopheles gambiae* and *Aedes aegypti*, these limitations have been overcome through the development of stable transgenic lines that express *Cas9* and *gRNA* endogenously, often coupled with dual visible markers to facilitate identification of edited progeny[8, 9]. This strategy depends critically on the availability of germline-biased promoters for *Cas9* and RNA polymerase III promoters for *gRNA* expression. The selection of species-specific promoters with high transcriptional activity is therefore essential for robust *invivo* expression of CRISPR components[10, 11]. Nevertheless, such well-characterized regulatory elements remain scarce for many non-model yet medically important mosquito species, including *Anopheles sinensis*, significantly impeding progress in functional genetics and targeted control strategies in these vectors.

Here, we developed and characterized germline-biased promoters and *U6* promoters capable of driving *Cas9*and *gRNA* expression in *Anopheles sinensis*. Using these regulatory elements, we established transgenic lines harboring stably integrated *Cas9* and *gRNA* expression cassettes and demonstrated that crosses between these lines achieve 100% germline editing efficiency, providing a robust and heritable genome editing platform for this neglected malaria vector.

Spatiotemporal expression profiling confirmed predominant transcription of *Asvasa2, Aszpg*, and *Asnanos* in female gonads (Figure 1A), consistent with the germline-biased patterns of their orthologs in other mosquito species[12, 13]. Using regulatory sequences from these genes, we generated transgenic constructs and established corresponding *Cas9*-expressing lines (Figure 1B, Figure S1, Table S1). The *Asvasa2-Cas9* line exhibited strong fluorescence in nurse cells by 48 h and throughout developing eggs at 70 h post-blood feeding, while signal intensity was remarkably reduced in *Aszpg-Cas9* and nearly absent in *Asnanos-Cas9* lines (Figure 1B, Figure S2). qPCR analysis further confirmed that the *Cas9* mRNA levels were substantially higher in the ovaries of *Asvasa2-Cas9* mosquitoes relative to the other two lines (Figure 1B). Notably, *Asvasa2-Cas9* also drove stronger *Cas9* expression in male testes relative to *Aszpg-Cas9* and *Asnanos-Cas9* (Figure S3).

**Figure 1.**
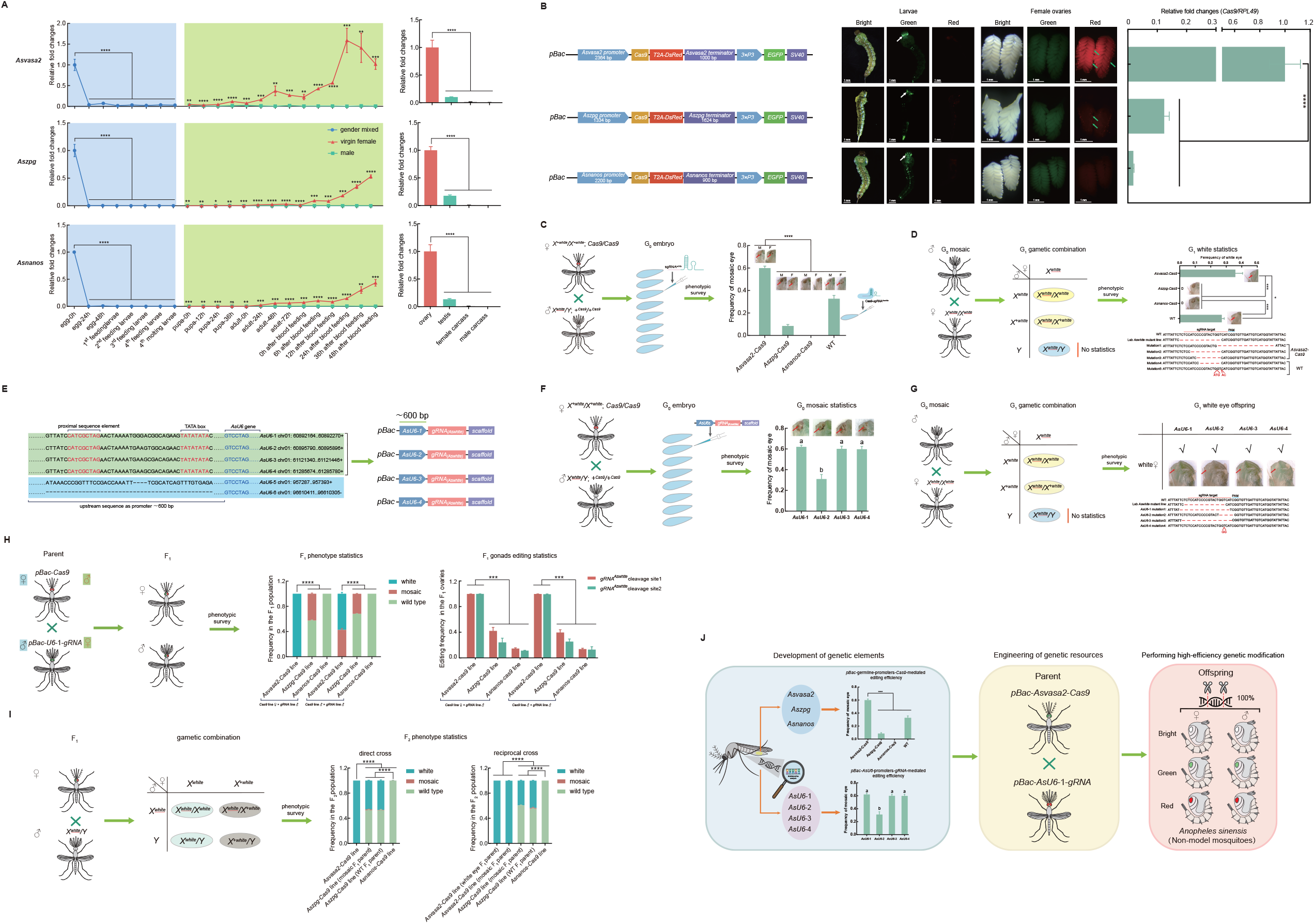
An engineered promoter system enables highly efficient transgenic-based genome editing in the malaria mosquito *Anophelessinensis*. A. Spatiotemporal transcription profiles of *Asvasa2, Aszpg*, and *Asnanos*. Blue and green shading denote sexually undifferentiated and differentiated stages, respectively (n = 3; ^*^*P*< 0.05, ^**^*P*< 0.01, ^***^*P*< 0.001, ^****^*P*< 0.0001). B. *Cas9* expression and fluorescence in ovaries of transgenic lines under control of germline-biased promoters. White and green arrows indicate green and red fluorescence, respectively (n = 3; ^****^*P*< 0.0001; scale bar: 1 mm). C. Editing efficiency in G_0_ progeny from crosses between *Cas9* transgenic females and lab-reared *Aswhite* mutant males, compared with conventional WT females (crossed with lab-reared *Aswhite* mutant males) following injection of Cas9-*sgRNA* ^*Aswhite*^ complexes (M: male; F: female; red arrows: mosaic eyes; n = 3; ^****^*P* < 0.0001). D. Germline transmission assay via G_0_ mosaic males crossed to lab-reared *Aswhite* mutant females. G_1_ mutations were verified by sequencing (n = 3; ^*^ *P*< 0.05, ^***^*P*< 0.001). E. Genomic features of *AsU6* promoter regions and gRNA expression cassette architecture. F. *Aswhite* editing efficiency mediated by four *AsU6-gRNA* plasmids in G_0_ individuals (Different letters indicate significant differences; n = 3; *P*< 0.0001). G. Qualitative assessment of germline transmission edits driven by four *AsU6* promoters. G_1_ mutations were sequence-verified. H. Phenotypic distribution (n = 3) and editing frequencies (n ≥ 2) at dual *gRNA*^*Aswhite*^ sites in F_1_ offspring from crosses between *Cas9* and *AsU6*-1*-gRNA* lines. ^***^*P* < 0.001. I. Phenotypic distribution of F2 progeny from crosses of F_1_ females (derived from *Cas9*× *AsU6*-1 crosses) to lab-reared *Aswhite* mutant males (n = 3; ^****^*P* < 0.0001). J. Schematic overview of the promoter-driven editing system development and implementation.

To determine editing efficacy driven by female germline-biased promoters, we observed that the *Asvasa2-Cas9* line induced mosaic phenotypes in 59.9% of G_0_ individuals—markedly higher than rates observed in WT (32.8%), *Aszpg-Cas9*(8.4%), and *Asnanos-Cas9*(0%) lines (Figure 1C, Table S2). Germline transmission was confirmed in G_1_ progeny, with 35.9% of females from the *Asvasa2-Cas9* group and 27.1% from WT microinjection group exhibiting the white-eyed phenotype (Figure 1D, Table S3, File S1). In contrast, no heritable edits were detected in G_1_ offspring derived from *Aszpg-Cas9* or *Asnanos-Cas9* G_0_ males (Figure 1D, Table S3). These results demonstrate that endogenous *Cas9* expression under germline-biased promoters provides a robust alternative to direct protein injection. The pronounced differences in efficiency among orthologous promoters underscores the importance of developing functional regulatory elements tailored to each target organism.

To identify functional Pol III promoters, we screened the *Anopheles sinensis* genome and identified six putative *AsU6* genes, four of which contained essential promoter elements—the proximal sequence element and TATA box (Figure 1E). Editing efficiency mediated by these promoters varied substantially: *AsU6*-2 yielded the lowest activity (31.1%), while *AsU6*-1 achieved the highest (62.1%), slightly exceeding *AsU6*-3 (60.2%) and *AsU6*-4 (59.9%) (Figure 1F, Table S4). All four *AsU6*promoters supported germline transmission, as evidenced by white-eyed G_1_ females following crosses of mosaic G_0_ males with lab-reared *Aswhite*mutant females (Figure 1G, File S2). Furthermore, when delivering the *AsU6*-1*-gRNA* construct was delivered into other two *Cas9* backgrounds, the *Aszpg-Cas9*line showed a mosaic editing rate of 10.3%, while no edits were detected in the *Asnanos-Cas9* background (Figure S4, Table S5). Although *AsU6*-3 and *AsU6*-4 exhibited activities comparable to *AsU6*-1, their suitability for germline editing requires further validation to enable diversified regulatory toolkits for gene drive applications and mitigate risks associated with repeated use of identical promoters[14].

Leveraging the validated promoters as a foundation, we next established a promoter-driven transgenic system to assess heritable editing efficiency. Direct crosses between *Asvasa2-Cas9* females and *AsU6*-1*-gRNA*^*Aswhite*^ males produced 100% white-eyed F_1_ progeny; while in the reciprocal cross, 57.4% of F_1_ offspring were fully white-eyed, while the remaining 42.6% exhibited mosaic eyes with ∼92.6% unpigmented area (Figure 1H, S5, S6; Table S1, S6). The complete absence of mosaic phenotype in the direct cross suggests that maternal Cas9 deposition from oocytes enhanced editing efficacy in early embryos[15]. Ovarian editing frequencies exceeded 99.3% in F_1_ females from both cross schemes (Figure 1H, Table S7, File S3). All F_2_ offspring from subsequent crosses of F_1_ females to lab-reared *Aswhite* mutant males displayed the white-eyed phenotype (Figure 1I, Table S8), demonstrating 100% germline transmission using the *Asvasa2*and *AsU6*-1 promoters.

Crosses between *Aszpg-Cas9* females and *AsU6-1-gRNA*^*Aswhite*^ males produced 42.6% F_1_ offspring with mosaic eyes (∼19.3% unpigmented area), while the reciprocal cross yielded 32.0% mosaics (∼8.3% unpigmented area) (Figure 1H, S5, S6; Table S6). Ovarian editing rates in F_1_ females ranged from 24.2% to 42.1% (Figure 1H, Table S7, File S3), and subsequent F_2_ crosses (obtained from from crosses of mosaic or wild-type F_1_ females with white-eyed males) produced white-eyed progeny at frequencies of 39.4–46.8% (Figure 1I, Table S9). These results indicate that the *Aszpg* promoter supports moderate germline editing, consistent with its utility in *Anopheles gambiae* gene-drive systems[13]. In contrast, crosses involving *Asnanos-Cas9* and *AsU6*-1*-gRNA* lines yielded no detectable eye phenotypes in F_1_ (Figure 1H, S5, S6; Table S6), low ovarian editing rates (≤ 14.6%; Figure 1H, Table S7, File S3), and minimal white-eyed F_2_ offspring (≤ 0.2%; Figure 1I, Table S10). Collectively, these data demonstrate that the *Asvasa2* promoter drives *Cas9* expression with superior efficiency and stability in the germline compared to *Aszpg* and *Asnanos*.

We also evaluated the *AsU6*-2 promoter in combination with *Asvasa2-Cas9*. Direct crosses produced 36.8% white-eyed and 63.2% mosaic F_1_ progeny (∼88.2% unpigmentation) (Figure S6, S7, Table S11). No white-eyed individuals occurred in the reciprocal cross, and mosaic eyes showed reduced unpigmentation (∼53.8% unpigmentation) (Figure S6–S8; Table S11). Ovarian editing rates ranged from 60.1% to 94.6% (Figure S9, Table S12, File S3), and white-eyed F_2_ frequencies were ≤ 62.5% (Figure S10, Table S13), and appreciably lower than those with *AsU6*-1, confirming that *AsU6*-2 drives functional but less efficient editing.

In summary, we have established a promoter-driven transgenic system that enables highly efficient gene editing in *Anopheles sinensis*, a major malaria vector (Figure 1J). Our findings greatly expands the genetic toolkit available for synthetic biology, such as gene drive-based population suppression, in non-model mosquitoes, as well as providing a transferable conceptual framework for genetic modification in other recalcitrant insect species.

## Acknowledgments

This work was supported by the National Natural Science Foundation of China (Nos. 31772527, 31872262); Chongqing Natural Science Foundation Innovation Development Joint Fund (CSTB2025NSCQ-LZX0119); the Scientific and Technological Research Program of Chongqing Municipal Education Commission (Nos. KJZD-K202200507; KJQN202200533; KJQN202200501); the Program of Bill & Melinda Gates Foundation (No. INV-061480); the Venture and Innovation Support Program for Chongqing Overseas Returnees (No. cx2022052); Chongqing “Express” Science and Research Program for PhD (No. CSTB2022BSXM-JCX0067); the Graduate Research and Innovation Foundation of Chongqing under Grant (No. CYS240374); and College Students’ Innovation and Entrepreneurship Training Plan Program (No. 202410637012).

## Author contributions

**Jun-Feng Hong**: Writing - Original draft, Writing - Review & Editing, Visualization, Methodology, Investigation, Data Curation. **Qi-Li Zou**: Writing - Review & Editing, Methodology, Validation, Investigation, Data curation. **Xin-Yuan Xie**: Methodology, Validation, Resources, Investigation, Data curation. **Yan-Ping Jiang**: Methodology, Validation, Investigation, Data curation. **Si-Yi Wang**: Investigation, Resources. **Xia Ling**: Methodology. **Cao Zhou**: Data Curation. **Wei Sun**: Data curation. **Xi Cai**: Resources. **Yong-Xin Yang**: Resources. **Yu Chen:** Resources. **Bin Chen**: Writing - Review & Editing, Supervision, Funding acquisition, Data curation, Conceptualization. **Liang Qiao**: Writing - Original draft, Writing - Review & Editing, Methodology, Validation, Supervision, Funding acquisition, Resources, Data curation, Conceptualization.

