## supplemental figures tables and files for "Engineered Promoter System Enables High-efficiency Transgenic CRISPR Editing in Malaria Transmitting Mosquito *Anopheles sinensis*": Supplemental figure 20251001 NEW.pdf

Figure S1

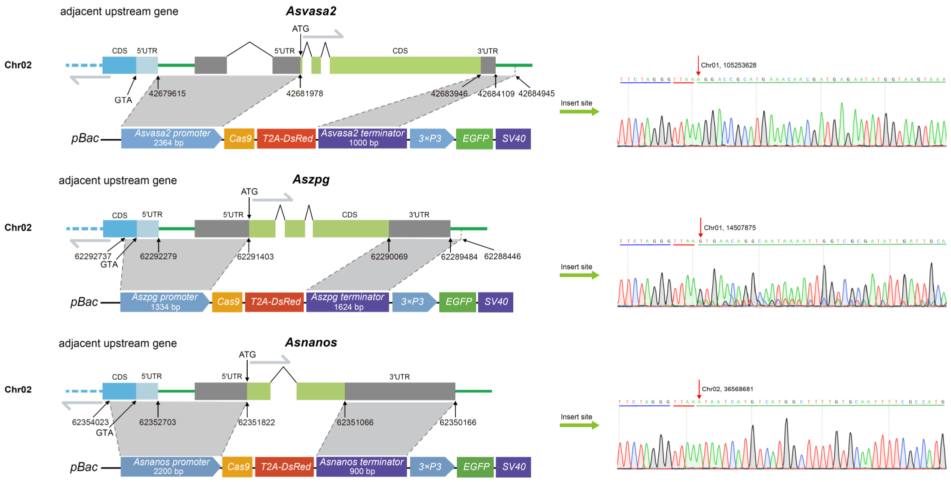


Schematic of promoter and terminator regions used for constructing the three *Cas9* expression plasmids and their genomic insertion sites in transgenic lines. Blue underlines denote the *piggyBac*-derived sequences, red underlines indicate the *TTAA* insertion site, and green underlines represent the flanking genomic regions.

Figure S2

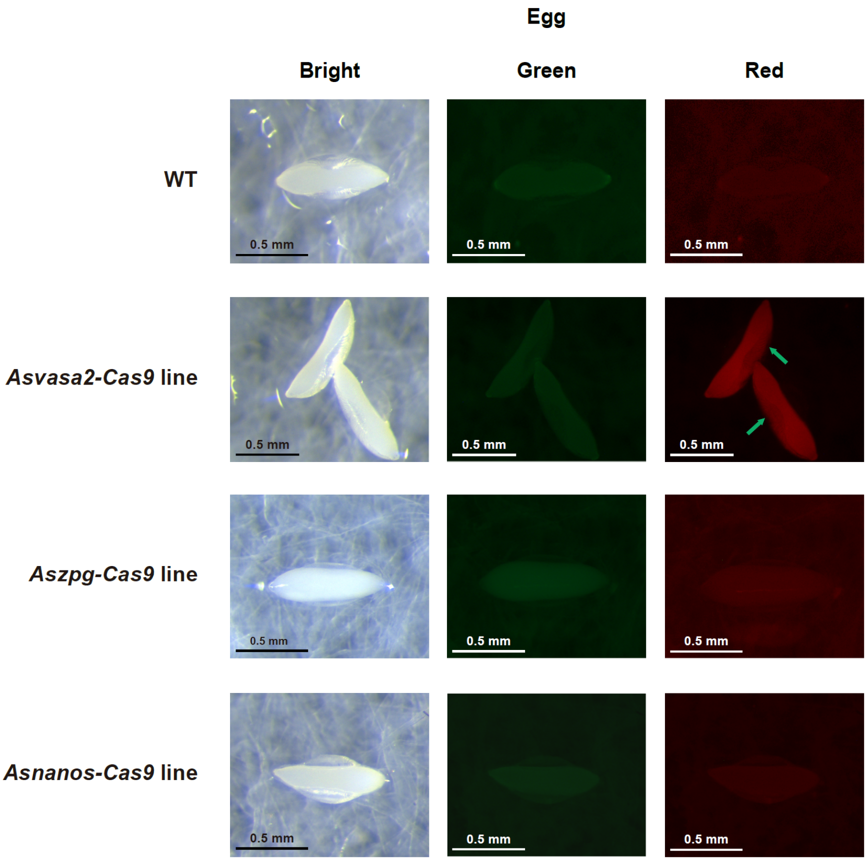


Fluorescent phenotypes of 0 h eggs laid by wild-type and three *Cas9* transgenic female lines at 70 h post-blood feeding. Green arrows highlight eggs displaying distinct red fluorescence.Scale bar = 0.5 mm.

Figure S3

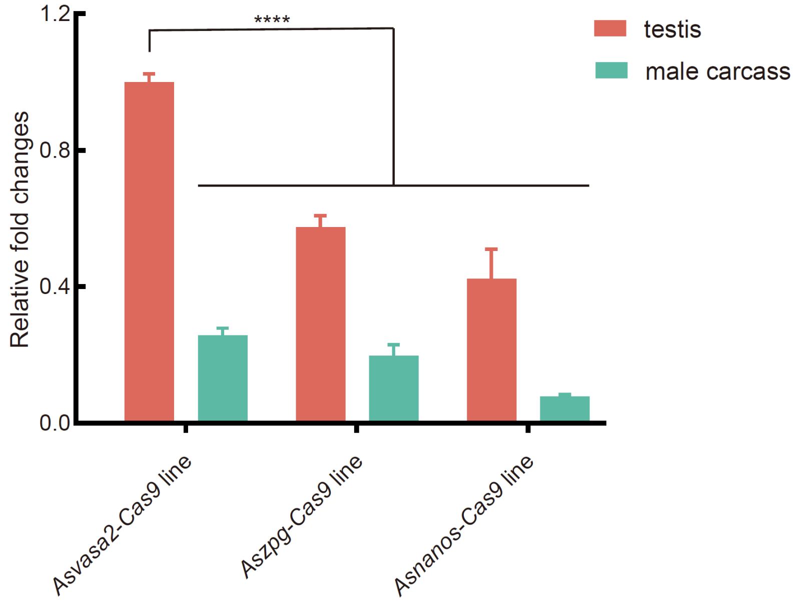


The *Cas9* transcripts levels in the testes and carcass of 2-day-old adult males from three *Cas9* transgenic lines (n = 3, ****$P$ < 0.0001).

Figure S4

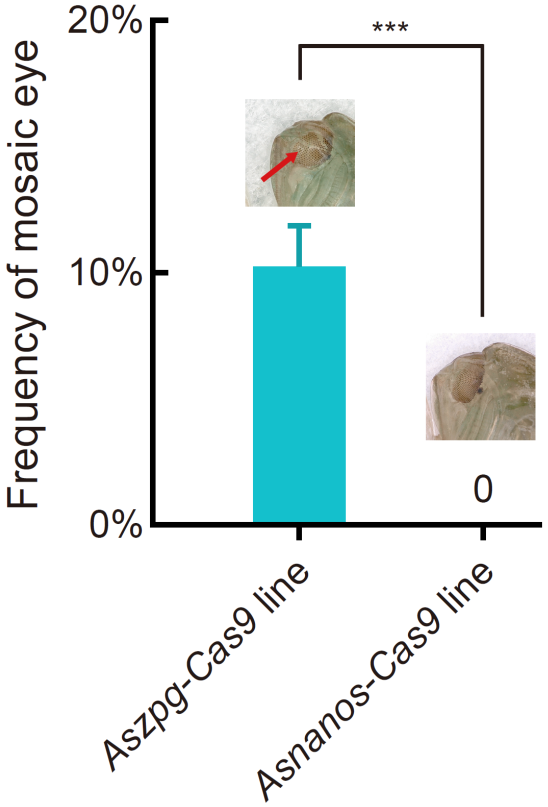


Mosaic-eyed phenotype frequencies in $F_1$ progeny following transient injection of the *AsU6*-1-*gRNA*$^{Aswhite}$ plasmid into embryos derived from crosses between *Aszpg*-*Cas9* or *Asnanos*-*Cas9* females and lab-reared *Aswhite* mutant males. The *Aszpg*-*Cas9* line represents progeny from *Aszpg*-*Cas9* females × lab-reared *Aswhite* mutant males, and the *Asnanos*-*Cas9* line represents progeny from *Asnanos*-*Cas9* females × lab-reared *Aswhite* mutant males, (n = 3, ****P* < 0.001).

Figure S5

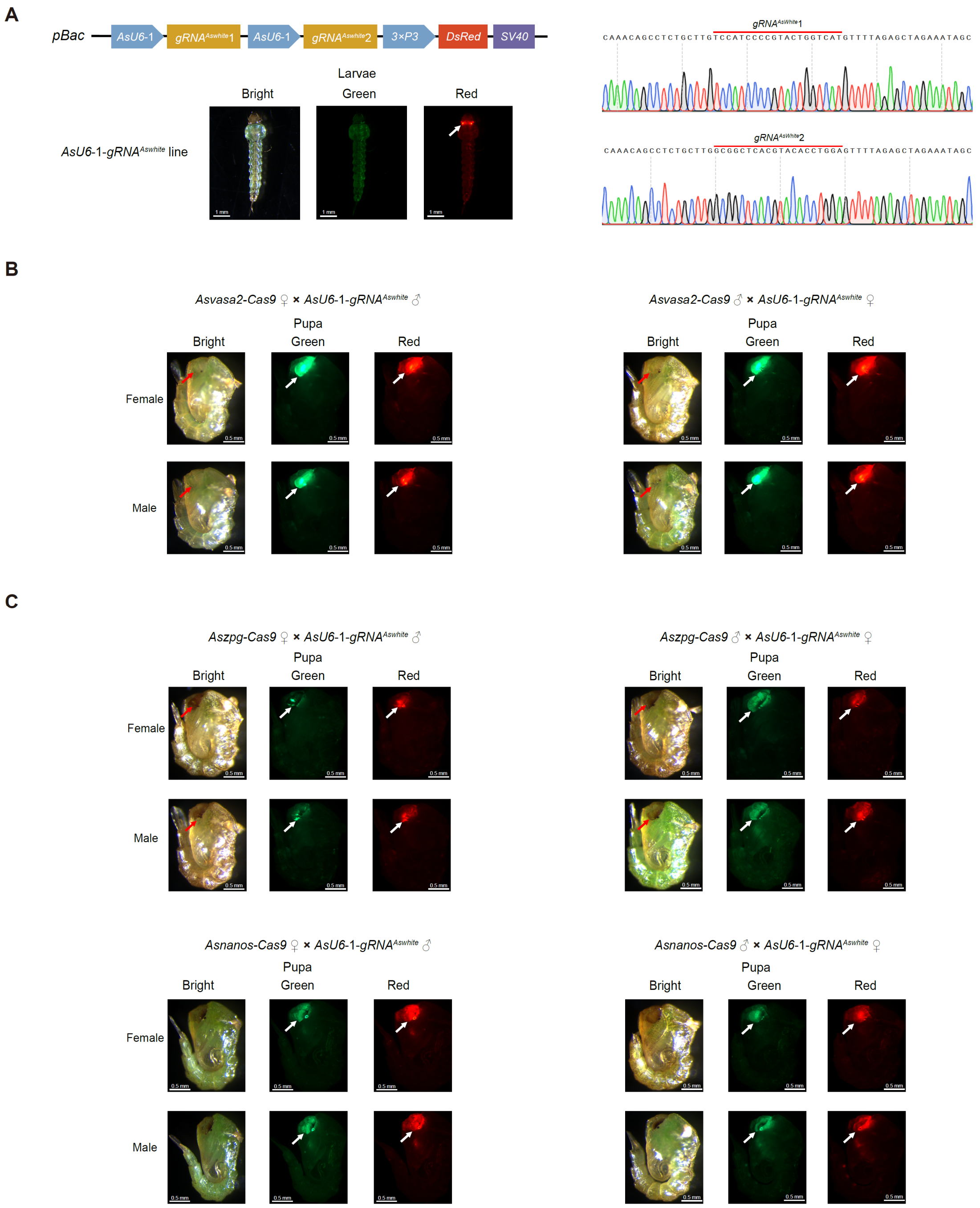


Generation of the *AsU6*-1-*gRNA*$^{Aswhite}$ line and its genetic crosses with *Cas9* transgenic lines. A. Plasmid schematic and sequencing chromatograms of the *gRNA* target sites used for construction of the *AsU6*-1-*gRNA*$^{Aswhite}$ line, together with the fluorescent phenotype of positive larvae. B. Eye and fluorescent phenotypes of $F_1$ pupae from crosses between the *AsU6*-1-*gRNA*$^{Aswhite}$ line and the *Asvasa*-*Cas9* line. C. Eye and fluorescent phenotypes of $F_1$ pupae from crosses with the *Aszpg*-*Cas9* or *Asnanos*-*Cas9* lines. White arrow indicates fluorescence; red arrow denotes mutant phenotype (white or mosaic eye).

Figure S6

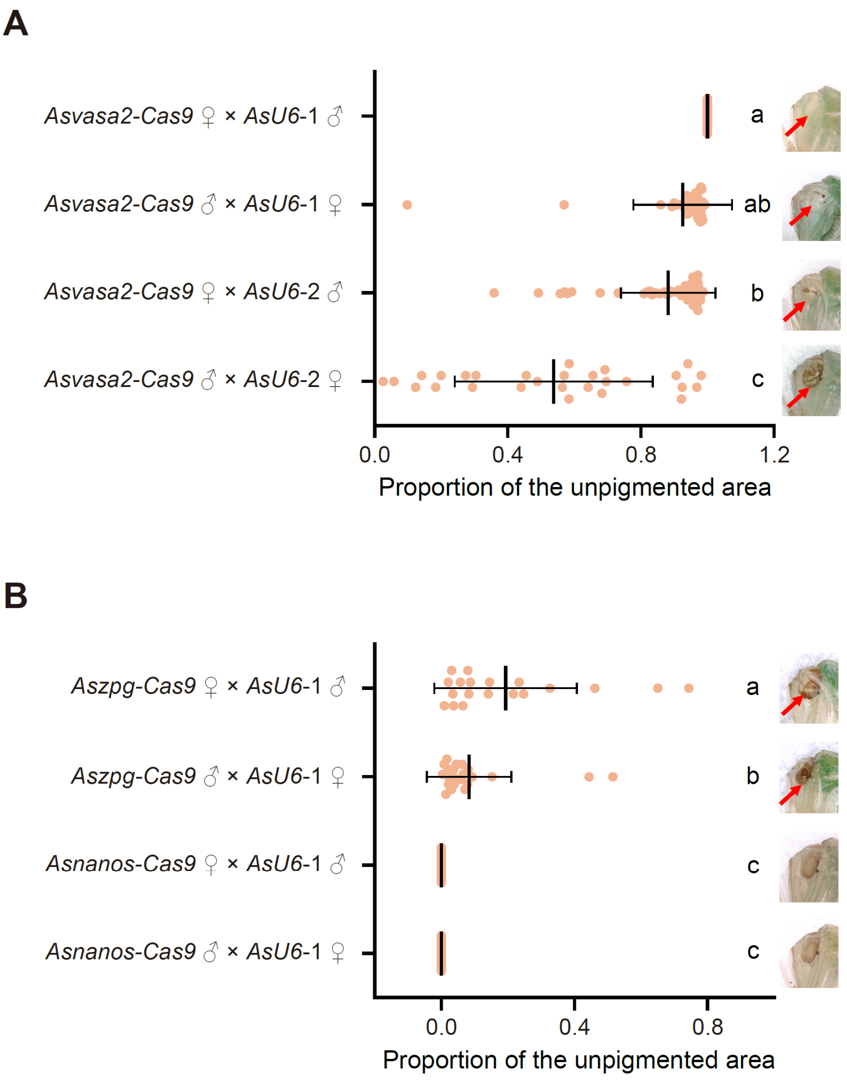


Proportion of the unpigmented area in the compound eye of pupal offspring derived from the '*Cas9* × *AsU6*' transgenic crosses. A. Ratios in $F_1$ pupae from hybrid crosses between the *Asvasa2*-*Cas9* line and the *AsU6*-1 or *AsU6*-2 line. B. Ratios in $F_1$ pupae from hybird crosses between the *Aszpg*-*Cas9* or *Asnanos*-*Cas9* line and the *AsU6*-1 line. (Different letters to the right of scatter plots indicate significant differences; n ≥ 19, $P < 0.05$), Red arrow indicates mutant phenotype (white or mosaic eye).

Figure S7

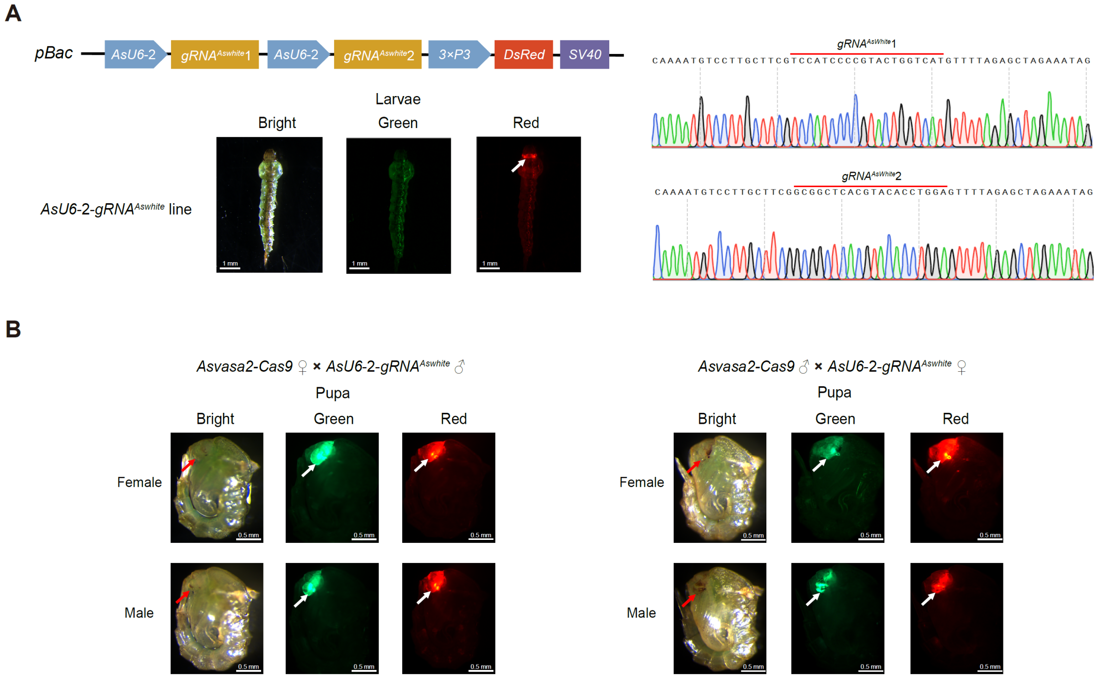


Construction of the *AsU6-2-gRNA*$^{Aswhite}$ line and its crosses with *Cas9* transgenic lines. A. Plasmid schematic and sequencing chromatograms of the *gRNA* target sites used to establish the *AsU6-2-gRNA*$^{Aswhite}$ line, together with the fluorescent phenotype of positive larvae. B. Eye and fluorescent phenotypes of $F_1$ pupa from crosses between the *AsU6-2-gRNA*$^{Aswhite}$ line and the *Asvasa-Cas9* line. White arrow indicates fluorescence; red arrow indicates mutant phenotype (white or mosaic eye).

Figure S8

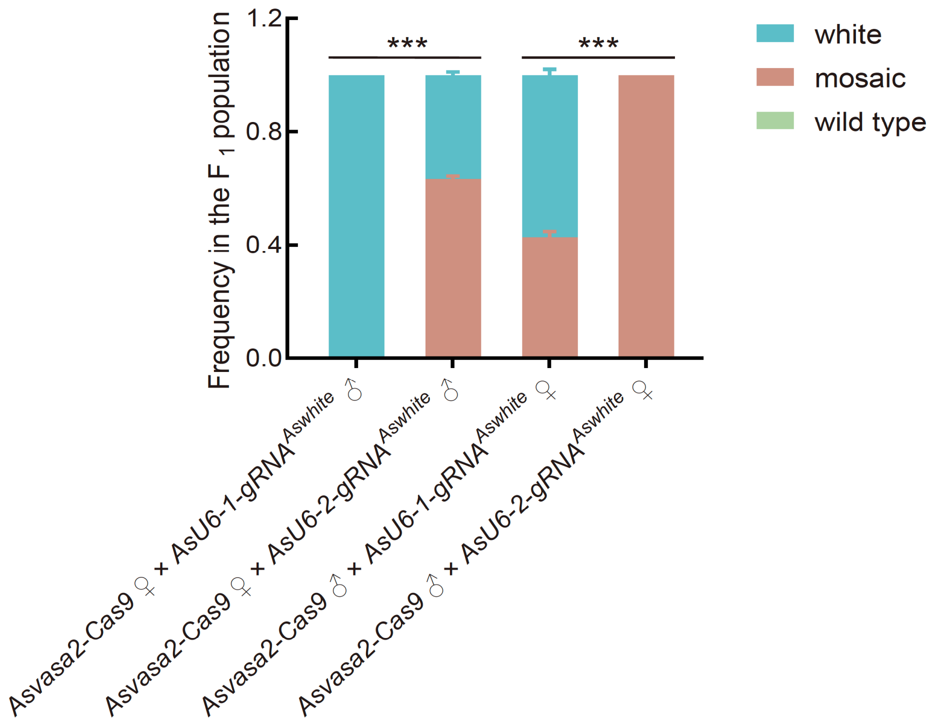


Phenotypic frequencies of white-eyed individuals in $F_1$ offspring from crosses between the *Asvasa2*-*Cas9* line and either the *AsU6*-1 or *AsU6*-2 lines. Statistical analyses of white-eyed phenotype proportions were performed with 3 biological repeats, ***$P < 0.0001$.

Figure S9

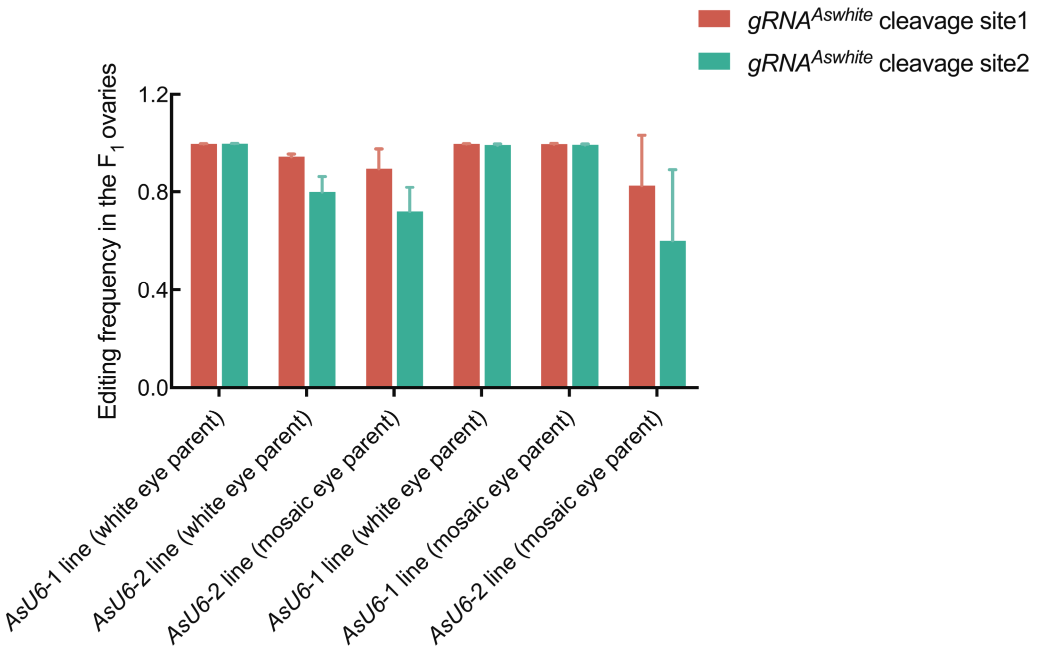


Editing efficiency at dual *gRNA*$^{Aswhite}$ target sites in ovarian tissues of white-eyed and mosaic $F_1$ females from crosses between the *Asvasa2*-*Cas9* line and either the *AsU6*-1 or *AsU6*-2 lines, (n ≥ 2).

Figure S10

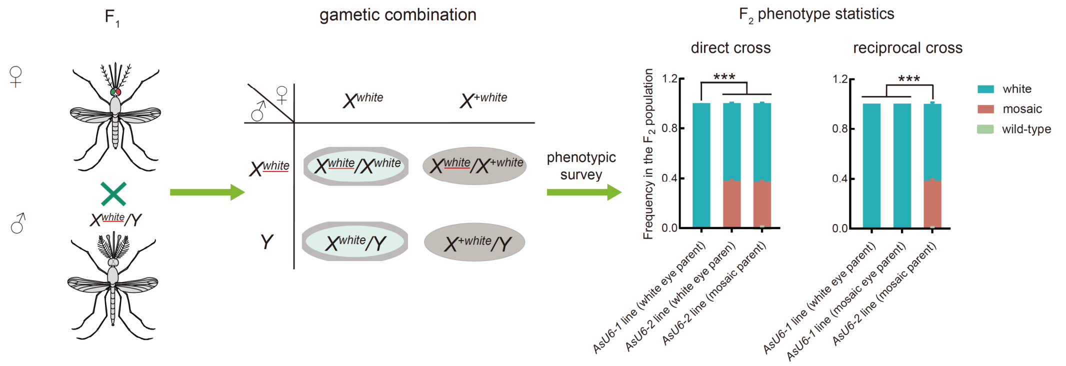


Phenotypic statistics of white-eyed individuals in $F_2$ offspring derived from crosses of $F_1$ females with distinct eye phenotypes (from hybrid crosses between the *Asvasa2*-*Cas9* and the *AsU6*-1/*AsU6*-2 lines) with lab-reared *Aswhite* mutant males. Statistical analyses of white-eyed phenotype proportions were performed with 3 biological repeats, ****P* < 0.001. Direct cross: $F_1$ individuals from *Asvasa2*-*Cas9* females × *AsU6* males; reciprocal cross: $F_1$ individuals from *Asvasa2*-*Cas9* males × *AsU6* females.
