## supplemental figures tables and files for "Engineered Promoter System Enables High-efficiency Transgenic CRISPR Editing in Malaria Transmitting Mosquito *Anopheles sinensis*": Supplemental files 20251001 NEW.pdf

File S1.Representative sequencing chromatograms showing mutation types in $G_1$ white-eyed females

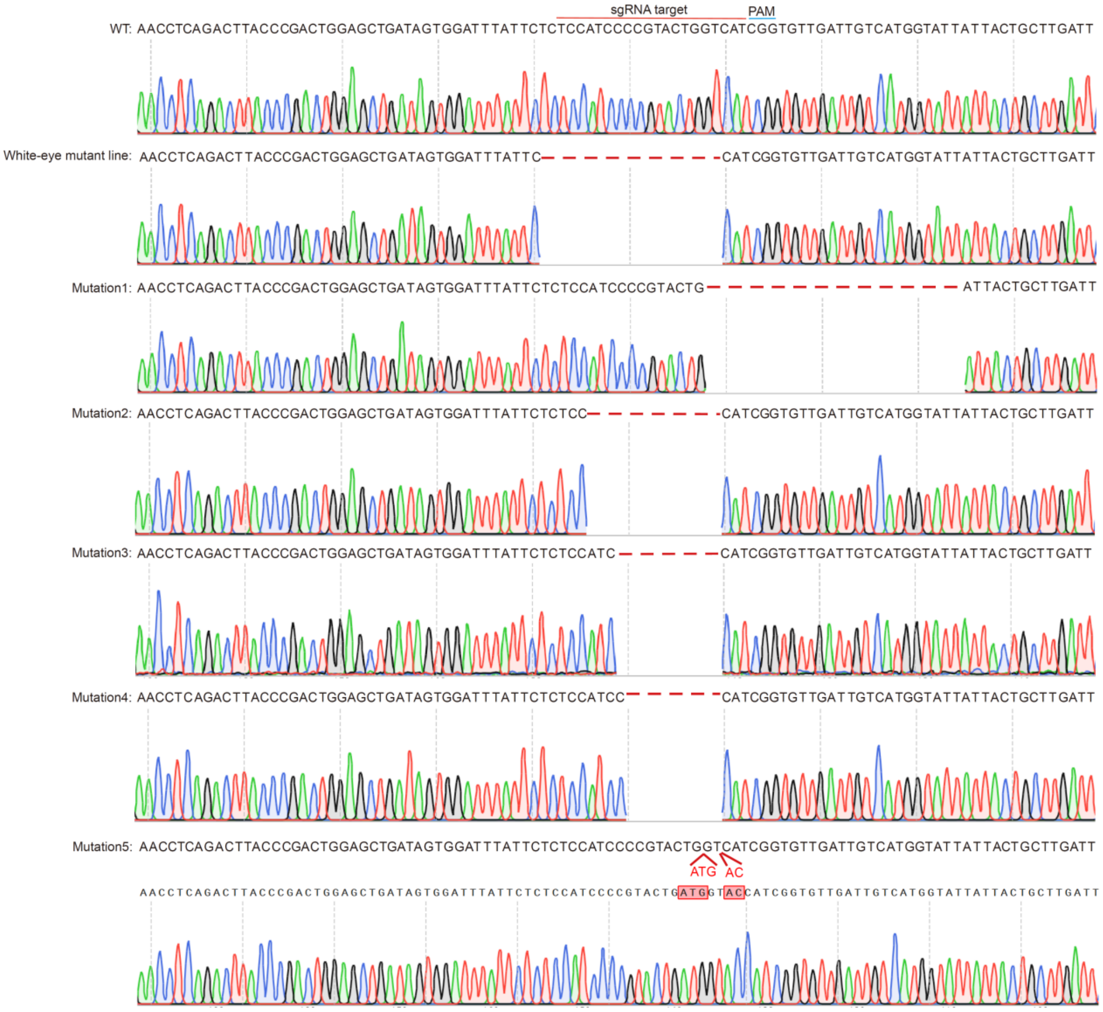


Confirmed mutagenesis in $G_1$ white-eyed females following crosses between *sgRNA*-injected $G_0$ mutant males (from *Cas9* or WT lines) and lab-reared *Aswhite* mutant females. Mutation 1-3 are from the *Asvasa2*-*Cas9* line, and mutation 4-5 are from the WT line. The top line indicates the wild-type sequence, the second line shows lab-reared *Aswhite* mutant line, and deletions or insertions are highlighted in red.

File S2. Representative sequencing chromatograms showing mutation types in $G_1$ white-eyed females

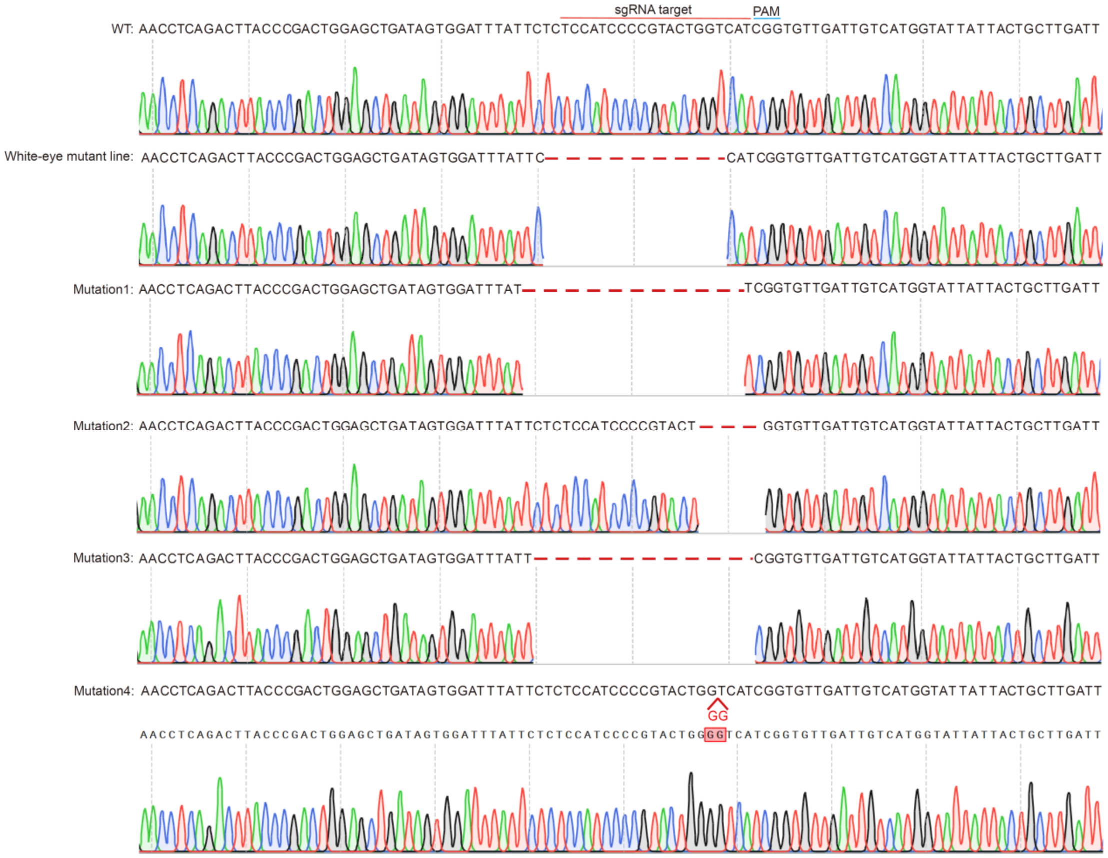


Confirmed mutagenesis in $G_1$ white-eyed females following crosses between $G_0$ mutant males (generated by injection of four *AsU6-gRNA*$^{Aswhite}$ plasmids into the *Asvasa2-Cas9* line) and lab-reared *Aswhite* mutant females. Mutations 1-4 correspond to *AsU6*-1-*gRNA*$^{Aswhite}$ to *AsU6*-4-*gRNA*$^{Aswhite}$ groups. The top line represents the wild-type sequence, the second line shows the lab-reared *Aswhite* mutant line, and deletions or insertions are highlighted in red.

File S3. Top 10 mutation types in $F_1$ ovaries from *Cas9* × As*U6* crosses, ranked by editing event frequencies and identified via high-throughput sequencing

a. Edit events at *gRNA*$^{Aswhite}$ sites in $F_1$ ovaries (white eye $F_1$ from *Asvasa2-Cas9* line ♀ × *AsU6-1-gRNA*$^{Aswhite}$ ♂)

| | | *gRNA*$^{Aswhite1}$ | | *gRNA*$^{Aswhite2}$ | |
|---|---|---|---|---|---|
| | SeqName | Sequence features | Frequency | Sequence features | Frequency |
| individual 1 | wild type | ACAATCAACACCGATGACCAGTACGGGGATGGAGAGAATAAAT | 0.28% | CGCAGCACGAGCGGCTCACGTACACCTGGAAGGAGATCGACGT | 0.09% |
|  | mutation 1 | ACAATCAACACCGATG--------GGGGATGGAGAGAATAAAT | 26.07% | CGCAGCACGAGCGGCTC----------GGAAGGAGATCGACGT | 27.79% |
|  | mutation 2 | ACAATCAACACCGATGTCCAGTACGAACACCGAACACCAGTACGGGGATGGAGAGAATAAAT | 8.28% | CGCAGCACGAGCGG-----------TGGAAGGAGATCGACGT | 18.81% |
|  | mutation 3 | ACAATCAACACCGATGA-------------------------- | 8.00% | CGCAGCACGAGCGGCTCACGTACACCTCACGGCTCACGGAAGGAGATCGACGT | 9.98% |
|  | mutation 4 | ACAATCAACACCGAT-----GTACGGGGATGGAGAGAATAAAT | 5.98% | CGCAGCACGAGCGG-----------CTGGAAGGAGATCGACGT | 6.56% |
|  | mutation 5 | ACAATCAACACCGAACATTACCAGTACGTAATACGGGGATGGAGAGAATAAAT | 5.07% | CGCAGCACGAGCGGCT---------CTGGAAGGAGATCGACGT | 5.55% |
|  | mutation 6 | ACAATCAACACCGATGA-------GGGGATGGAGAGAATAAAT | 4.03% | CGCAGCACGAGCGGCTCACGTA-----GGAAGGAGATCGACGT | 3.23% |
|  | mutation 7 | ACAATCAACACCGATG-CCAGTACGGGGATGGAGAGAATAAAT | 4.02% | CGCAGCACGAGCGGCTCAC--------GGAAGGAGATCGACGT | 2.54% |
| | mutation 8 | ACAATCAACACCGATACATTCACCGATCGTCCAGTACGGGTCGAACCGATCGTACCAGTACGGGGATGGAGAGAATAAAT | 3.86% | CGCAGCACGAGCGGCTCACGTACACCTTGGAAGGAGATCGACGT | 2.52% |
|  | mutation 9 | ACAATCAACACCGATGA---------GGATGGAGAGAATAAAT | 3.63% | CGCAGCACGAGCGGCTCAC---------GAAGGAGATCGACGT | 1.95% |
|  | mutation 10 | ACAATCAACACCGATG-------CGGGGATGGAGAGAATAAAT | 3.40% | CGCAGCACGAGCGGCTCACGTACACCGAACACGGAAGGAGATCGACGT | 1.87% |
| individual 2 | wild type | ACAATCAACACCGATGACCAGTACGGGGATGGAGAGAATAAAT | 0.17% | CGCAGCACGAGCGGCTCACGTACACCTGGAAGGAGATCGACGT | 0.18% |
|  | mutation 1 | ACAATTG--------GATCA--ACGGGGATGGAGAGAATAAAT | 40.09% | CGCAGCACGAGCGGCTCACGTACACCTTGGAAGGAGATCGACGT | 12.79% |
|  | mutation 2 | ACAATCAACACCGAT-----------GGATGGAGAGAATAAAT | 7.61% | CGCAGCACGAGCGGCTC----------GGAAGGAGATCGACGT | 9.35% |
|  | mutation 3 | ACAATCAACACCGATG--------GGGGATGGAGAGAATAAAT | 6.33% | CGCAGCACGAGCGG-------------GGAAGGAGATCGACGT | 8.09% |
|  | mutation 4 | ACAATCAACACCGAT----------GGGATGGAGAGAATAAAT | 6.04% | CGCAGCACGAGCGG-----------CTGGAAGGAGATCGACGT | 7.33% |
|  | mutation 5 | ACAATCAACACCGAT------------GATGGAGAGAATAAAT | 4.40% | CGCAGCACGAGCGGC------------GGAAGGAGATCGACGT | 4.99% |
| | mutation 6 | ACAATCAACACCGATGAACCAGTACGGGGATGGAGAGAATAAAT | 3.52% | CGCAGCACGAGCGGCTCACGTACACCGTACACGTACGGAAGGAGATCGACGT | 4.66% |
|  | mutation 7 | ACAATCAACACC---GACCAGTACGGGGATGGAGAGAATAAAT | 3.06% | CGCAGCACGAGCGGCTCAC--------GGAAGGAGATCGACGT | 4.65% |
|  | mutation 8 | ACAATCAACACCGAT-----GTACGGGGATGGAGAGAATAAAT | 2.19% | CGCAGCACGAGCGGCT----------TGGAAGGAGATCGACGT | 4.39% |
|  | mutation 9 | ACAATCAACACCGATG------ACGGGGATGGAGAGAATAAAT | 2.09% | CGCAGCACGA----------------TGGAAGGAGATCGACGT | 4.38% |
|  | mutation 10 | ACAATCAACACCGAT---------GGGGATGGAGAGAATAAAT | 1.67% | CGCAGCA------------------CTGGAAGGAGATCGACGT | 4.27% |

b. Edit events at *gRNA*$^{Aswhite}$ sites in $F_1$ ovaries (white eye $F_1$ from *Asvasa2-Cas9* line ♂ × *AsU6-1-gRNA*$^{Aswhite}$ ♀)

| | | *gRNA*$^{Aswhite1}$ | | *gRNA*$^{Aswhite2}$ | |
|---|---|---|---|---|---|
| | SeqName | Sequence features | Frequency | Sequence features | Frequency |
| individual 1 | wild type | ACAATCAACACCGATGACCAGTACGGGGATGGAGAGAATAAAT | 0.34% | CGCAGCACGAGCGGCTCACGTACACCTGGAAGGAGATCGACGT | 0.99% |
|  | mutation 1 | ACAATCAAC-------ACCAGTACGGGGATGGAGAGAATAAAT | 20.81% | CGCAGCACGAGCGGCTCACGTACACCGACGGAAGGAGATCGACGT | 19.09% |
|  | mutation 2 | ACAATCAACACCGATG--------GGGGATGGAGAGAATAAAT | 8.06% | CGCAGCACGAGCGGCTCAC--------GGAAGGAGATCGACGT | 10.22% |
|  | mutation 3 | ACAATCAACACCGAT-----------GGATGGAGAGAATAAAT | 6.54% | CGCAGCACGAGCGGCTC----------GGAAGGAGATCGACGT | 7.39% |
|  | mutation 4 | ACAATCAACACCGAT---------GGGGATGGAGAGAATAAAT | 6.38% | CGCAGCACGAGCGGCT---------CTGGAAGGAGATCGACGT | 5.05% |
|  | mutation 5 | ACAATCAACACCGATG------------------------AAAT | 4.87% | CGCAGCACGAGCGGCTCACGTACACCTTGGAAGGAGATCGACGT | 4.14% |
|  | mutation 6 | ACAATCAACACCGAT----------GGGATGGAGAGAATAAAT | 4.49% | CGCAGCACGAGCGGCTCACG-------GGAAGGAGATCGACGT | 3.35% |
|  | mutation 7 | ACAATCAACACCGATG------ACGGGGATGGAGAGAATAAAT | 4.26% | --------------------------TGGAAGGAGATCGACGT | 3.19% |
|  | mutation 8 | ACAATCAACACCGATG--------------------------- | 4.12% | CGCAGCACGAGCGG-------------GGAAGGAGATCGACGT | 3.11% |
|  | mutation 9 | ACAATCAACACCGAT------------GATGGAGAGAATAAAT | 3.65% | CGCAGC---------------------GGAAGGAGATCGACGT | 2.57% |
|  | mutation 10 | ACAATCAACACCGA-------TACGGGGATGGAGAGAATAAAT | 2.56% | CGCAGCACGAGCGG-----------CTGGAAGGAGATCGACGT | 2.00% |
| individual 2 | wild type | ACAATCAACACCGATGACCAGTACGGGGATGGAGAGAATAAAT | 0.20% | CGCAGCACGAGCGGCTCACGTACACCTGGAAGGAGATCGACGT | 0.41% |
|  | mutation 1 | ACAATCAACACCGATG--------GGGGATGGAGAGAATAAAT | 20.36% | CGCAGCACGA-----------------GGAAGGAGATCGACGT | 19.42% |
|  | mutation 2 | ACAATCAACACCGATG------ACGGGGATGGAGAGAATAAAT | 11.51% | CGCAGCACGAGCGGCTC----------GGAAGGAGATCGACGT | 17.21% |
|  | mutation 3 | ACAATCAACACCGAT-----------GGATGGAGAGAATAAAT | 11.24% | CGCAGCACGAGCGGCTCACGTACACCTTGGAAGGAGATCGACGT | 8.85% |
|  | mutation 4 | ACAATCAAC-------ACCAGTACGGGGATGGAGAGAATAAAT | 11.10% | CGCAGCACGAGCGG-----------CTGGAAGGAGATCGACGT | 8.76% |
|  | mutation 5 | ACAATCAACACC---------------------AGAGAATAAAT | 8.43% | CGCAGCACGAG---------------TGGAAGGAGATCGACGT | 5.46% |
|  | mutation 6 | ACAATCAACACCGAT-----------------GAGAGAATAAAT | 7.37% | CGCAGCACGAGCGGCTCACGTACACCTCTGGAAGGAGATCGACGT | 4.33% |
|  | mutation 7 | ACAATCAACACCGAT---------GGGGATGGAGAGAATAAAT | 5.92% | CGCAGCACGAGCGGCTCACGT----CTGGAAGGAGATCGACGT | 4.23% |
|  | mutation 8 | ACAATCAACACCGATG----GTACGGGGATGGAGAGAATAAAT | 5.61% | CGCAGCACGAGCGGCTCAC--------GGAAGGAGATCGACGT | 4.17% |
|  | mutation 9 | ACAATCAACACCGAT---------------------GAATAAAT | 2.59% | CGCAGCACGAGCGGCTCACG------TGGAAGGAGATCGACGT | 3.52% |
| | mutation 10 | ACAATCAACACCGATCATGATCATGATCGGGGATGGAGAGAATAAAT | 2.28% | CGCAGCACGAGCGGCTCACGTACACCT-GAAGGAGATCGACGT | 2.12% |

c. Edit events at *gRNA*$^{Aswhite}$ sites in $F_1$ ovaries (mosaic eye $F_1$ from *Asvasa2-Cas9* line ♂ × *AsU6-1-gRNA*$^{Aswhite}$ ♀)

| | | *gRNA*$^{Aswhite1}$ | | *gRNA*$^{Aswhite2}$ | |
|---|---|---|---|---|---|
| | SeqName | Sequence features | Frequency | Sequence feature | Frequency |
| individual 1 | wild type | ACAATCAACACCGATGACCAGTACGGGGATGGAGAGAATAAAT | 0.16% | CGCAGCACGAGCGGCTCACGTACACCTGGAAGGAGATCGACGT | 0.37% |
|  | mutation1 | ACAATCAACACCGAT-----------GGATGGAGAGAATAAAT | 19.51% | CGCAGCACGAGCGGCTCACGTACACCGAAGGAAGGAGATCGACGT | 17.75% |
|  | mutation2 | ACAATCAAC-------ACCAGTACGGGGATGGAGAGAATAAAT | 14.54% | CGCAGCACGAGAGG-----------TTGGAAGGAGATCGACGT | 11.22% |
|  | mutation3 | ACAATCAACACCGAT------------GATGGAGAGAATAAAT | 12.27% | CGCAGCACGAGCGG-------------GGAAGGAGATCGACGT | 10.97% |
|  | mutation4 | ACAATCAACACCGAT---------GGGGATGGAGAGAATAAAT | 11.91% | CGCAGCACGAGCGGCT---------CTGGAAGGAGATCGACGT | 10.64% |
|  | mutation5 | ACAATCAACACCGATG-------CTGGGATGGAGAGAATAAAT | 8.61% | CGCAGCACGAGCGGCTCAC---------GAAGGAGATCGACGT | 10.61% |
|  | mutation6 | ACAATCAACACCGAT-----GTACGGGGATGGAGAGAATAAAT | 8.16% | CGCAGCACGAGCGGCTCA--------TGGAAGGAGATCGACGT | 10.18% |
|  | mutation7 | ACAATCAACACCGAT-----------------GAGAGAATAAAT | 7.18% | CGCAGCACGAGCG-------------TGGAAGGAGATCGACGT | 8.11% |
|  | mutation8 | ACAATCAACACCGAT----------GGGATGGAGAGAATAAAT | 6.57% | CGCAGCACGAGCGGC------------GGAAGGAGATCGACGT | 4.90% |
|  | mutation9 | ACAATCAACACCGATG-------------TGGAGAGAATAAAT | 0.47% | CGCAGCACGA----------------TGGAAGGAGATCGACGT | 4.38% |
|  | mutation10 | ACAATCAACACCGATG------ACGGGGATGGAGAGAATAAAT | 0.40% | CGCAGCACGA---------------CTGGAAGGAGATCGACGT | 1.23% |
| individual 2 | wild type | ACAATCAACACCGATGACCAGTACGGGGATGGAGAGAATAAAT | 0.51% | CGCAGCACGAGCGGCTCACGTACACCTGGAAGGAGATCGACGT | 0.98% |
|  | mutation1 | ACAATCAACACCGATG--------GGGGATGGAGAGAATAAAT | 17.64% | CGCAGCACGAGCGG-----------CTGGAAGGAGATCGACGT | 12.09% |
|  | mutation2 | ACAATCAACACCGAT------------GATGGAGAGAATAAAT | 11.26% | CGCAGCACGAGCGGCTCACGTACACCTTGGAAGGAGATCGACGT | 8.90% |
|  | mutation3 | ACAATCAACACCGAT-----------GGATGGAGAGAATAAAT | 6.58% | CGCAGCACGAG---------------TGGAAGGAGATCGACGT | 8.89% |
|  | mutation4 | ACAATCAACACCGATG------ACGGGGATGGAGAGAATAAAT | 6.53% | CGCAGCACGAGCGGCTC----------GGAAGGAGATCGACGT | 8.60% |
|  | mutation5 | ACAATCAACACCGATG----------------------ATAAAT | 5.95% | CGCAGCACGAGCGGCTCAC--------GGAAGGAGATCGACGT | 8.00% |
|  | mutation6 | ACAATCAACACCGATG-------CGGGGATGGAGAGAATAAAT | 5.66% | CGCAGCACGAGCGG------------TGGAAGGAGATCGACGT | 4.67% |
|  | mutation7 | ACAATCAACACCGAT----------GGGATGGAGAGAATAAAT | 4.78% | CGCAGCACGAGCGGCT----------TGGAAGGAGATCGACGT | 4.66% |
|  | mutation8 | ACAATCAACACCGATGA--------------------AATAAAT | 4.37% | CGCAGCACGA----------------TGGAAGGAGATCGACGT | 4.11% |
|  | mutation9 | ACAATCAACACCGAT-----GTACGGGGATGGAGAGAATAAAT | 3.94% | CGCAGCACGAGCGGC------------GGAAGGAGATCGACGT | 3.95% |
|  | mutation10 | ACAATCAACACCGATG-----------------------TAAAT | 3.70% | CGCAGCACGAGCGG-------------GGAAGGAGATCGACGT | 3.79% |

d. Edit events at *gRNA*$^{Aswhite}$ sites in $F_1$ ovaries (mosaic eye $F_1$ from *Aszpg-Cas9* line ♀ × *AsU6-1-gRNA*$^{Aswhite}$ ♂)

| | | $gRNA^{Aswhite1}$ | | $gRNA^{Aswhite2}$ | |
|---|---|---|---|---|---|
| | SeqName | Sequence features | Frequency | Sequence features | Frequency |
| individual 1 | wild type | ACAATCAACACCGATGACCAGTACGGGGATGGAGAGAATAAAT | 59.75% | CGCAGCACGAGCGGCTCACGTACACCTGGAAGGAGATCGACGT | 77.38% |
|  | mutation 1 | ACAATCAACACCGAT-ACCAGTACGGGGATGGAGAGAATAAAT | 6.63% | CGCTCCGCAAGCGA-----AAAAAAAGGGAAGGAGATCGACGT | 4.70% |
|  | mutation 2 | ACAATCAACACCGATG--------GGGGATGGAGAGAATAAAT | 2.36% | CGCAGCACGAGCGGCTCACGTACACC-GGAAGGAGATCGACGT | 1.38% |
| | mutation 3 | ACAATCAACACCGATCAACCAGTACGGGGATGGAGAGAATAAAT | 1.82% | CGCAGCACGAGCGGCTCACGTACAC--GGAAGGAGATCGACGT | 0.68% |
| | mutation 4 | ACAATCAACACCGATG-CCAGTACGGGGATGGAGAGAATAAAT | 1.39% | CGCAGCACGAGCGGCTCACGTACACGCAGCGGGAAGGAGATCGACGT | 0.66% |
|  | mutation 5 | ACAATCAAC-------ACCAGTACGGGGATGGAGAGAATAAAT | 1.39% | CGCAGCACGAGCGGCCCACGTACACCTGGAAGGAGATCGACGT | 0.40% |
| | mutation 6 | ACAATCAACACC---GACCAGTACGGGGATGGAGAGAATAAAT | 1.25% | CGCAGCACGAGCGGCTCACGTACACCTGGAAGGGGATCGACGT | 0.36% |
| | mutation 7 | ACAATCAACACCGATGGGGATGGAGTACCAGTACGGGGATGGAGAGAATAAAT | 1.10% | CGCAGCACGAGCGGCTCACGTACGCCTGGAAGGAGATCGACGT | 0.34% |
| | mutation 8 | ACAATCAACACCGATGGTACCAGTACGGGGATGGAGAGAATAAAT | 1.00% | CGCAGCACGAGCGGCTCACGTACACCTGGAGGGAGATCGACGT | 0.33% |
|  | mutation 9 | ACAATCAACACCGAT-----GTACGGGGATGGAGAGAATAAAT | 0.93% | CGCAGCACGAGCGGCTCACGTACACCTGGAAGGAGGTCGACGT | 0.33% |
| | mutation 10 | ACAATCAACACCGATGGTGTACGTACGGGGATGGAGAGAATAAAT | 0.87% | CGCAGCACGAGCGGCTCACGTACACGTGTCGTACAAGGAGATCGACGT | 0.32% |
| individual 2 | wild type | ACAATCAACACCGATGACCAGTACGGGGATGGAGAGAATAAAT | 54.63% | CGCAGCACGAGCGGCTCACGTACACCTGGAAGGAGATCGACGT | 66.16% |
|  | mutation 1 | ACAATCAACACCGAT----------GGATGGAGAGAATAAAT | 13.90% | CGCAGCACGAGCGGCTCACGTAC----GGAAGGAGATCGACGT | 9.07% |
| | mutation 2 | ACAATCAACACCGATGTACGGGATGTACCAGTACGGGGATGGAGAGAATAAAT | 9.07% | CGCAGCACGAGCGGCTCACGTACAC--GGAAGGAGATCGACGT | 7.03% |
|  | mutation 3 | ACAATCAACACCGATG--------GGGGATGGAGAGAATAAAT | 2.51% | CGCAGCACGAGCGGCTCACGTACACGTACACGTACACGGAAGGAGATCGACGT | 1.39% |
|  | mutation 4 | ACAATCAACACCGAT----------------GAGAGAATAAAT | 1.28% | CGCAGCACGAGCGGCTCACGTACACC-GGAAGGAGATCGACGT | 0.48% |
| | mutation 5 | ACAATCAACAGCGATGACCAGTACGGGGATGGAGAGAATAAAT | 0.76% | CGCAGCACGAGCGGCTCACGTACACGAGCGGAAGGAGATCGACGT | 0.44% |
|  | mutation 6 | ACAATCAACACCGAT------------------AGAATAAAT | 0.46% | CGCAGCACGAGCGGCTCACGTACACCTGGAAGGAGGTCGACGT | 0.38% |
|  | mutation 7 | ACAATCAACACCGATGTGAACCCCGACCAGTACGGGGATGGAGAGAATAAAT | 0.44% | CGCAGCACGAGCGGAGA----------GGAAGGAGATCGACGT | 0.37% |
| | mutation 8 | ACAATCAACACCGATGAACCAGTACGGGGATGGAGAGAATAAAT | 0.40% | CGCAGCACGGGCGGCTCACGTACACCTGGAAGGAGATCGACGT | 0.35% |
|  | mutation 9 | ACAATCAACACCGATG------ACGGGGATGGAGAGAATAAAT | 0.34% | CGCAGCACGAGCGGCTCACGTACACCTGGAAGGAGATCGGCGT | 0.34% |
|  | mutation 10 | ACAATCAACACCGAT---------GGGGATGGAGAGAATAAAT | 0.33% | CGCAGCACGAGCGGCTCGCGTACACCTGGAAGGAGATCGACGT | 0.34% |

e. Edit events at $gRNA^{Aswhite}$ sites in $F_1$ ovaries (wild-type eye $F_1$ from *Aszpg-Cas9* line ♀ × *AsU6-1-gRNA*$^{Aswhite}$ ♂ )

| | | $gRNA^{Aswhite1}$ | | $gRNA^{Aswhite2}$ | |
|---|---|---|---|---|---|
| | SeqName | Sequence feature | Frequency | Sequence features | Frequency |
| individual 1 | wild type | ACAATCAACACCGATGACCAGTACGGGGATGGAGAGAATAAAT | 64.57% | CGCAGCACGAGCGGCTCACGTACACCTGGAAGGAGATCGACGT | 80.03% |
| | mutation 1 | ACAATCAACATGGGGATGGACAACCAGTACGGGGATGGAGAGAATAAAT | 3.44% | CGCAGCACGAGCGGCTCACGATCAC--GGAAGGAGATCGACGT | 1.47% |
|  | mutation 2 | ACAATCAACACCGATGG--AGTACGGGGATGGAGAGAATAAAT | 3.39% | CGCAGCACGAGCGGC-----------GGAAGGAGATCGACGT | 1.09% |
|  | mutation 3 | ACAATCAAC-------ACCAGTACGGGGATGGAGAGAATAAAT | 2.13% | CGCAGCACGAGCGGCTCACGTACACCTCACGGAAGGAGATCGACG | 1.01% |
|  | mutation 4 | ACAATCAACACCGATG------ACGGGGATGGAGAGAATAAAT | 1.45% | CGCAGCACGAGCGGCTCACGTACGCGCGGCTCACGCGGGAAGGAGATCGACGT | 0.90% |
|  | mutation 5 | ACAATCAACACCGATG--------GGGGATGGAGAGAATAAAT | 1.36% | CGCAGCACGAGCGGCTCACGTACACCGAAGGAAGGAAGGAAGGAGATCGACGT | 0.78% |
|  | mutation 6 | ACAATCAACACCG---------TCGGGGATGGAGAGAATAAAT | 1.36% | CGCAGCACGAGCGGCCCACGTACACCTGGAAGGAGATCGACGT | 0.43% |
|  | mutation 7 | ACAATCAACACCGAT----------GGATGGAGAGAATAAAT | 1.21% | CGCAGCACGAGCGGCTCACGTACACCTGGAAGGAGGTCGACGT | 0.42% |
|  | mutation 8 | ACAATCA---------GCCAGTACGGGGATGGAGAGAATAAAT | 1.17% | CGCAGCACGAGCGGCTCACGTACACC-GGAAGGAGATCGACGT | 0.40% |
| | mutation 9 | ACAATCAACACCGATCAACACCAGTACGGGGATGGAGAGAATAAA | 1.02% | CGCAGCACGGGCGGCTCACGTACACCTGGAAGGAGATCGACGT | 0.38% |
| | mutation 10 | ACAATCAACATC---GACCAGTACGGGGATGGAGAGAATAAAT | 1.00% | CGCAGCACGAGCGGCTCACGTACACCTGGAGGGAGATCGACGT | 0.36% |
| individual 2 | wild type | ACAATCAACACCGATGACCAGTACGGGGATGGAGAGAATAAAT | 52.47% | CGCAGCACGAGCGGCTCACGTACACCTGGAAGGAGATCGACGT | 79.67% |
|  | mutation 1 | ACAATCAAC-------ACCAGTACGGGGATGGAGAGAATAAAT | 7.63% | CGCAGCACGAGCGGCTCACGTACAC---GAAGGAGATCGACGT | 1.69% |
|  | mutation 2 | ACAATCAACACCGATG-------GGGGATGGAGAGAATAAAT | 3.63% | CGCAGCACGAGCGGCTCACG------TGGAAGGAGATCGACGT | 1.57% |
|  | mutation 3 | ACAATCAACACCGAT----------GGATGGAGAGAATAAAT | 2.88% | CGCAGCACGAGC--------------GGAAGGAGATCGACGT | 1.23% |
|  | mutation 4 | ACAAT------------CAGTACGGGGATGGAGAGAATAAAT | 2.57% | CGCAGCACGAGCGGCTCACGTACAC--GGAAGGAGATCGACGT | 0.85% |
| | mutation 5 | ACAATCAACACCGATG--CAGTACGGGGATGGAGAGAATAAAT | 2.33% | CGCAGCACGAGCGGCTCACGTGAGTTCGACGTC-GGAAGGAGATCGACGT | 0.73% |
|  | mutation 6 | ACAATCAACACCGATG----GTACGGGGATGGAGAGAATAAAT | 1.75% | CGCAGCACGAGCGGCT---------CTGGAAGGAGATCGACGT | 0.46% |
|  | mutation 7 | ACAATCAAC-------TCCAGTACGGGGATGGAGAGAATAAAT | 1.70% | CGCAGCACGAGCGGCTCACGTACACCTACAGCGTACGTGCAGCGTGCGTACCTACGTACGTACAGCGGAAGGAGATCGACGT | 0.42% |
|  | mutation 8 | ------TACGT-----ACTAGTACGGGGATGGAGAGAATAAAT | 1.37% | CGCAGCACGAGCGGCTCACGTACACCTGGAAGGAGGTCGACGT | 0.39% |
| | mutation 9 | ACAATCAACACCGTACGTAGATCCCAGTACGGGGATGGAGAGAATAAAT | 1.16% | CGCAGCACGAGCGGCTCACGTACACCTGGAAGGGGATCGACGT | 0.38% |
|  | mutation 10 | ACAATCAACACCGAT---------GGGATGGAGAGAATAAAT | 1.04% | CGCAGCACGAGCGGCCCACGTACACCTGGAAGGAGATCGACGT | 0.36% |

f. Edit events at $gRNA^{Aswhite}$ sites in $F_1$ ovaries (mosaic eye $F_1$ from *Aszpg-Cas9* line ♂ × *AsU6-1-gRNA*$^{Aswhite}$ ♀ )

| | | $gRNA^{Aswhite1}$ | | $gRNA^{Aswhite2}$ | |
|---|---|---|---|---|---|
| | SeqName | Sequence features | Frequency | Sequence features | Frequency |
| individual 1 | wild type | ACAATCAACACCGATGACCAGTACGGGGATGGAGAGAATAAAT | 63.68% | CGCAGCACGAGCGGCTCACGTACACCTGGAAGGAGATCGACGT | 73.80% |
|  | mutation 1 | ACAATCAAC-------ACCAGTACGGGGATGGAGAGAATAAAT | 12.12% | CGCAGCACGAGCGGCTCACGTACACCACGGAAGGAGATCGACGT | 5.80% |
|  | mutation 2 | ACAATCAACACCGATG--------GGGGATGGAGAGAATAAAT | 1.95% | CGCAGCACGAGCGGCTCACGTACTCCTTCTAAGGAGATCGACGT | 5.61% |
|  | mutation 3 | ACAATCAACACCGAT-----------GATGGAGAGAATAAAT | 1.53% | CGCAGCACGAGCGGCTCACGTACACCTTGGAAGGAGATCGACGT | 0.91% |
|  | mutation 4 | ACAATCAACACCGATGAC--------GGATGGAGAGAATAAAT | 1.02% | CGCAGCACGAGCGGCTCAC--------GGAAGGAGATCGACGT | 0.87% |
|  | mutation 5 | ACAATCAACACCGAT----------GGATGGAGAGAATAAAT | 1.02% | CGCAGCACGAGCGGCCCACGTACACCTGGAAGGAGATCGACGT | 0.39% |
|  | mutation 6 | ACAATCAACACCGAT-----GTACGGGGATGGAGAGAATAAAT | 0.80% | CGCAGCACGAGCGGCTCACGTACACCTGGACGGAGATCGACGT | 0.38% |
| | mutation 7 | ACAATCAACACCGATG--CAGTACGGGGATGGAGAGAATAAAT | 0.71% | CGCAGCACGAGCGGCTCACGTACACCTGGAAGGAGGTCGACGT | 0.36% |
|  | mutation 8 | ACAATCAACACCGAT---------GGGATGGAGAGAATAAAT | 0.69% | CGCAGCACGAGCGGCTCACGTACACCTGGAGGGAGATCGACGT | 0.36% |
| | mutation 9 | ACAATCAACACCGATG-CCAGTACGGGGATGGAGAGAATAAAT | 0.60% | CGCAGCACGAGCGGCTCACGTACACCTGGAAGGGGATCGACGT | 0.34% |
|  | mutation 10 | ACAATCAACACCGAT-------------------GAATAAAT | 0.48% | CGCAGCACGAGCGGCTCACGTACGCCTGGAAGGAGATCGACGT | 0.33% |
| individual 2 | wild type | ACAATCAACACCGATGACCAGTACGGGGATGGAGAGAATAAAT | 63.27% | CGCAGCACGAGCGGCTCACGTACACCTGGAAGGAGATCGACGT | 78.13% |
|  | mutation 1 | ACAATCAACACCGAT----------GGATGGAGAGAATAAAT | 3.31% | CGCAGCACGAGCGGCTCACGTACACCTGGACGGAGATCGACGT | 4.14% |
|  | mutation 2 | ACAATCAAC-------ACCAGTACGGGGATGGAGAGAATAAAT | 2.80% | CGCAGCACGAGCG-------------GGAAGGAGATCGACGT | 0.96% |
|  | mutation 3 | ACAATCAACACCGATG--------GGGGATGGAGAGAATAAAT | 2.41% | CGCAGCACGAGCGGCTC----------GGAAGGAGATCGACGT | 0.85% |
|  | mutation 4 | ACAATCAACACCGAT---------GGGATGGAGAGAATAAAT | 2.34% | CGCAGCACGAGCGGCTCAC--------GGAAGGAGATCGACGT | 0.41% |
|  | mutation 5 | ACAATCAACACCGAT-----------GATGGAGAGAATAAAT | 1.54% | CGCAGCACGAGCGGCTCACGT--------AAGGAGATCGACGT | 0.39% |
|  | mutation 6 | ACAATCAACACCGATG----GTACGGGGATGGAGAGAATAAAT | 1.19% | CGCAGCACGAGCGGCTCACGTACACCTGGAGGGAGATCGACGT | 0.39% |
|  | mutation 7 | ACAATCAACACC----------------------GAATAAAT | 0.84% | CGCAGCACGAGCGGCTCACGTACACCTGGAAGGGGATCGACGT | 0.39% |
|  | mutation 8 | ACAATCAACACCGAT-----GTACGGGGATGGAGAGAATAAAT | 0.82% | CGCAGCACGAGCGGCTCACGTACACCTGGAAGGAGGTCGACGT | 0.37% |
|  | mutation 9 | ACAATCAACACCGAT---------GGGGATGGAGAGAATAAAT | 0.81% | CGCAGCACGAGCGGCTCACGTACACC-GGAAGGAGATCGACGT | 0.36% |
| | mutation 10 | ACAATCAACACCGATG---AGTACGGGGATGGAGAGAATAAAT | 0.59% | CGCAGCACGAGCGGCCCACGTACACCTGGAAGGAGATCGACGT | 0.35% |

g. Edit events at $gRNA^{Aswhite}$ sites in $F_1$ ovaries (wild-type eye $F_1$ from *Aszpg-Cas9* line ♂ × *AsU6-1-gRNA*$^{Aswhite}$ ♀ )

| | | $gRNA^{Aswhite1}$ | | $gRNA^{Aswhite2}$ | |
|---|---|---|---|---|---|
| | SeqName | Sequence features | Frequency | Sequence features | Frequency |
| individual 1 | wild type | ACAATCAACACCGATGACCAGTACGGGGATGGAGAGAATAAAT | 60.22% | CGCAGCACGAGCGGCTCACGTACACCTGGAAGGAGATCGACGT | 77.07% |
|  | mutation1 | ACAATCAAC-------ACCAGTACGGGGATGGAGAGAATAAAT | 11.59% | CGCAGCACGAGCGGCTCACGTACACGGCTCACGGAAGGAGATCGACGT | 2.48% |
|  | mutation2 | ACAATCAACACCGATAGTACCGATCGACCAGTACGGGGATGGAGAGAATAAAT | 9.89% | CGCAGCACGAGCGGCTCAC--------GGAAGGAGATCGACGT | 2.34% |
|  | mutation3 | ACAATCAACACCGATG----GTACGGGGATGGAGAGAATAAAT | 0.86% | CGCAGCACGAGCGGCTCACGTACACC-GGAAGGAGATCGACGT | 1.25% |
|  | mutation4 | ACAATCAACACCGATG--------GGGGATGGAGAGAATAAAT | 0.71% | CGCAGCACGAGCGGCTCACGTACAC--GGAAGGAGATCGACGT | 0.73% |
|  | mutation5 | ACAATCAACACCGAT---------GGGGATGGAGAGAATAAAT | 0.41% | CGCAGCACGAGCGGCTCACGTAC----GGAAGGAGATCGACGT | 0.43% |
| | mutation6 | ACAATCAACACCGATGACCAGTACGGGGATGGAGAGGATAAAT | 0.35% | CGCAGCACGAGCGGCCCACGTACACCTGGAAGGAGATCGACGT | 0.42% |
| | mutation7 | ACAATCAACACCGATGAACCAGTACGGGGATGGAGAGAATAAAT | 0.35% | CGCAGCACGAGCGGCTCACGTACACCTGGAAGGAGGTCGACGT | 0.42% |
| | mutation8 | ACAATCAACACCGATGACCAGTACGGGGATGGGGAGAATAAAT | 0.34% | CGCAGCACGAGCGGCTCACGTACACACACGAAGGAGATCGACGT | 0.39% |
| | mutation9 | ACAATCAACACCGATGACCAGTACGGGGATGGAGGGAATAAAT | 0.30% | CGCAGCACGAGCGGCTCACGTACACCTGGAGGGAGATCGACGT | 0.37% |
| | mutation10 | GCAATCAACACCGATGACCAGTACGGGGATGGAGAGAATAAAT | 0.29% | CGCAGCACGAGCGGCTCACGTACGCCTGGAAGGAGATCGACGT | 0.35% |
| individual 2 | wild type | ACAATCAACACCGATGACCAGTACGGGGATGGAGAGAATAAAT | 54.44% | CGCAGCACGAGCGGCTCACGTACACCTGGAAGGAGATCGACGT | 69.22% |
|  | mutation1 | ACAATCAACACC---------------GATGGAGAGAATAAAT | 6.80% | CGCAGCACGAGCGGCTCACGTACACCGAGGAAGGAGATCGACGT | 3.58% |
|  | mutation2 | ACAATCAAC-------ACCAGTACGGGGATGGAGAGAATAAAT | 5.35% | CGCAGCACGAGCGGCTCACGTACACGTACGGGAAGGAGATCGACGT | 3.22% |
|  | mutation3 | ACAATCAACACC-----------------------GAATAAAT | 5.04% | CGCAGCACGAGCGGCTCACGTACAA--GGAAGGAGATCGACGT | 3.12% |
| | mutation4 | ACAATCAACACCGATGCCCAGTACGGGGATGGAGAGAATAAAT | 2.83% | CGCAGCACGAGCGGCTCACGTACAC--GGAAGGAGATCGACGT | 2.44% |
| | mutation5 | ACAATCAACACCGATGTTACCATATCAACATACATAACAGTACGGGGATGGAGAGAATAAAT | 2.37% | CGCAGCACGAGCGGCTCACGTACACCATCGTGTACGGAAGGAGATCGACGT | 1.60% |
| | mutation6 | ACAATCAACACCGATGACCCCAGTACGGGGATGGAGAGAATAAAT | 1.42% | CGCAGCACGAGCGGCTCACGTACACCGCTCGGGAAGGAGATCGACGT | 1.53% |
| | mutation7 | ACAATCAACACCGATGGTACCATCAGTACGGGGATGGAGAGAATAAAT | 1.21% | CGCAGCACGAGCGGCTCACGTACACCGTGAGGAAGGAGATCGACGT | 1.25% |
| | mutation8 | ACAATCAACACCGGGAATGA-CAGTACGGGGATGGAGAGAATAAAT | 0.86% | CGCAGCACGAGCGGCTCACGTACA-CGGGAAGGAGATCGACGT | 0.45% |
|  | mutation9 | ACAATCAACACCGAT-----------GGATGGAGAGAATAAAT | 0.76% | CGCAGCACGAGCGGCTCACGTACACCTGGAAGGAGATCGGCGT | 0.35% |
| | mutation10 | ACAATCAACACCGATG---AGTACGGGGATGGAGAGAATAAAT | 0.65% | CGCAGCACGAGCGGCCCACGTACACCTGGAAGGAGATCGACGT | 0.35% |

h. Edit events at $gRNA^{Aswhite}$ sites in $F_1$ ovaries (wild-type eye $F_1$ from *Asnanos-Cas9* line ♀ × *AsU6-1-gRNA*$^{Aswhite}$ ♂)

| | | $gRNA^{Aswhite1}$ | | $gRNA^{Aswhite2}$ | |
|---|---|---|---|---|---|
| | SeqName | Sequence features | Frequency | Sequence features | Frequency |
| individual 1 | wild type | ACAATCAACACCGATGACCAGTACGGGGATGGAGAGAATAAAT | 86.21% | CGCAGCACGAGCGGCTCACGTACACCTGGAAGGAGATCGACGT | 88.97% |
| | mutation 1 | ACAATCAACACCGATGACCAGTACGGGGATGGGGAGAATAAAT | 0.47% | CGCAGCACGAGCGGCCCACGTACACCTGGAAGGAGATCGACGT | 0.45% |
| | mutation 2 | ACAATCAACACCGATGACCAGTACGGGGATGGAGGGAATAAAT | 0.45% | CGCAGCACGAGCGGCTCACGTACACCTGGAAGGGGATCGACGT | 0.43% |
| | mutation 3 | ACAATCAACACCGATGACCAGTACGGGGATGGAGAGGATAAAT | 0.44% | CGCAGCACGAGCGGCTCACGTACACCTGGAAGGAGGTCGACGT | 0.40% |
| | mutation 4 | ACAATCAGCACCGATGACCAGTACGGGGATGGAGAGAATAAAT | 0.42% | CGCAGCACGAGCGGCTCACGTACACCTGGAGGGAGATCGACGT | 0.38% |
| | mutation 5 | ACAATCAACACCGATGGCCAGTACGGGGATGGAGAGAATAAAT | 0.41% | CGCAGCACGGGCGGCTCACGTACACCTGGAAGGAGATCGACGT | 0.37% |
| | mutation 6 | GCAATCAACACCGATGACCAGTACGGGGATGGAGAGAATAAAT | 0.40% | CGCAGCACGAGCGGCTCACGTACGCCTGGAAGGAGATCGACGT | 0.35% |
| | mutation 7 | ACAATCAACACCGATGACCAGTACGGGGATGGAGAGAATAAAC | 0.37% | CGCAGCACGAGCGGCTCACGTACACCTGGAAGGAGATCGGCGT | 0.35% |
| | mutation 8 | ACAATCAACGCCGATGACCAGTACGGGGATGGAGAGAATAAAT | 0.37% | CGCAGCACGAGCGGCTCGCGTACACCTGGAAGGAGATCGACGT | 0.34% |
| | mutation 9 | ACGATCAACACCGATGACCAGTACGGGGATGGAGAGAATAAAT | 0.36% | CGCAGCGCGAGCGGCTCACGTACACCTGGAAGGAGATCGACGT | 0.34% |
| | mutation 10 | ACAATCAACACCGATGACCAGTACGGGGGTGGAGAGAATAAAT | 0.34% | CGCAGCACGAGCGGCTCACGTACACCTGGAAGGAGATCGACGC | 0.32% |
| individual 2 | wild type | ACAATCAACACCGATGACCAGTACGGGGATGGAGAGAATAAAT | 84.64% | CGCAGCACGAGCGGCTCACGTACACCTGGAAGGAGATCGACGT | 88.35% |
| | mutation 1 | ACAATCAACAGCGATGACCAGTACGGGGATGGAGAGAATAAAT | 1.28% | CGCAGCACGAGCGGCTCACGTACACCTGGAAGGAGGTCGACGT | 0.47% |
| | mutation 2 | ACAATCAACACCGATGACCAGTACGGGGATGGGGAGAATAAAT | 0.51% | CGCAGCACGAGCGGCTCACGTACACCTGGAAGGGGATCGACGT | 0.45% |
| | mutation 3 | GCAATCAACACCGATGACCAGTACGGGGATGGAGAGAATAAAT | 0.44% | CGCAGCACGAGCGGCCCACGTACACCTGGAAGGAGATCGACGT | 0.42% |
| | mutation 4 | ACAATCAACACCGATGACCAGTACGGGGATGGAGAGGATAAAT | 0.44% | CGCAGCACGGGCGGCTCACGTACACCTGGAAGGAGATCGACGT | 0.42% |
| | mutation 5 | ACAATCAACACCGATGGCCAGTACGGGGATGGAGAGAATAAAT | 0.39% | CGCAGCACGAGCGGCTCACGTACACCTGGAGGGAGATCGACGT | 0.41% |
| | mutation 6 | ACAATCAGCACCGATGACCAGTACGGGGATGGAGAGAATAAAT | 0.39% | CGCAGCACGAGCGGCTCGCGTACACCTGGAAGGAGATCGACGT | 0.40% |
| | mutation 7 | ACAATCAACACCGATGACCAGTACGGGGGTGGAGAGAATAAAT | 0.38% | CGCAGCACGAGCGGCTCACGTACACCTGGAAGGAGATCGACGC | 0.36% |
| | mutation 8 | ACAATCAACACCGATGACCAGTACGGGGATGGAGGGAATAAAT | 0.37% | CGCAGCGCGAGCGGCTCACGTACACCTGGAAGGAGATCGACGT | 0.35% |
| | mutation 9 | ACGATCAACACCGATGACCAGTACGGGGATGGAGAGAATAAAT | 0.36% | CGCAGCACGAGCGGCTCACGTACGCCTGGAAGGAGATCGACGT | 0.35% |
| | mutation 10 | ACAATCAACACCGATGACCAGTACGGGGATGGAGAGAGTAAAT | 0.36% | CGCGGCACGAGCGGCTCACGTACACCTGGAAGGAGATCGACGT | 0.33% |

i. Edit events at $gRNA^{Aswhite}$ sites in $F_1$ ovaries (wild-type eye $F_1$ from *Asnanos-Cas9* line ♂ × *AsU6-1-gRNA*$^{Aswhite}$ ♀)

| | | $gRNA^{Aswhite1}$ | | $gRNA^{Aswhite2}$ | |
|---|---|---|---|---|---|
| | SeqName | Sequence features | Frequency | Sequence features | Frequency |
| individual 1 | wild type | ACAATCAACACCGATGACCAGTACGGGGATGGAGAGAATAAAT | 87.27% | CGCAGCACGAGCGGCTCACGTACACCTGGAAGGAGATCGACGT | 90.21% |
|  | mutation1 | ACAATCAAC-------ACCAGTACGGGGATGGAGAGAATAAAT | 0.56% | CGCAGCACGAGCGGCCCACGTACACCTGGAAGGAGATCGACGT | 0.44% |
| | mutation2 | ACAATCAACACCGATGACCAGTACGGGGATGGGGAGAATAAAT | 0.46% | CGCAGCACGAGCGGCTCACGTACACCTGGAAGGAGGTCGACGT | 0.43% |
| | mutation3 | ACAATCAACACCGATGACCAGTACGGGGATGGAGAGGATAAAT | 0.43% | CGCAGCACGAGCGGCTCACGTACACCTGGAGGGAGATCGACGT | 0.42% |
| | mutation4 | ACAATCAACACCGATGACCAGTACGGGGATGGAGGGAATAAAT | 0.42% | CGCAGCACGAGCGGCTCACGTACACCTGGAAGGGGATCGACGT | 0.40% |
| | mutation5 | ACA-TCAACACCGATGACCAGTACGGGGATGGAGAGAATAAAT | 0.41% | CGCAGCACGAGCGGCTCACGTACACCTGGAAGGAGATCGGCGT | 0.38% |
| | mutation6 | ACAATCAGCACCGATGACCAGTACGGGGATGGAGAGAATAAAT | 0.40% | CGCAGCACGAGCGGCTCACGTACGCCTGGAAGGAGATCGACGT | 0.37% |
| | mutation7 | ACAATCAACACCGATGGCCAGTACGGGGATGGAGAGAATAAAT | 0.38% | CGCAGCACGGGCGGCTCACGTACACCTGGAAGGAGATCGACGT | 0.36% |
| | mutation8 | ACAATCAACGCCGATGACCAGTACGGGGATGGAGAGAATAAAT | 0.38% | CGCAGCGCGAGCGGCTCACGTACACCTGGAAGGAGATCGACGT | 0.35% |
| | mutation9 | ACAATCAACACCGATGACCAGTACGGGGGTGGAGAGAATAAAT | 0.36% | CGCAGCACGAGCGGCTCGCGTACACCTGGAAGGAGATCGACGT | 0.35% |
| | mutation10 | ACAATCAACACCGATGACCAGTACGGGGATGGAGAGAATAAAC | 0.33% | CGCGGCACGAGCGGCTCACGTACACCTGGAAGGAGATCGACGT | 0.35% |
| individual 2 | wild type | ACAATCAACACCGATGACCAGTACGGGGATGGAGAGAATAAAT | 85.38% | CGCAGCACGAGCGGCTCACGTACACCTGGAAGGAGATCGACGT | 87.83% |
| | mutation1 | ACAATCAACACCGATGACCAGTACGGGGATGGAGAGGATAAAT | 0.48% | CGCAGCACGAGCGGCTCACGTACACCTGGAAGGAGGTCGACGT | 0.47% |
| | mutation2 | ACAATCAGCACCGATGACCAGTACGGGGATGGAGAGAATAAAT | 0.42% | CGCAGCACGAGCGGCCCACGTACACCTGGAAGGAGATCGACGT | 0.46% |
| | mutation3 | ACAATCAACACCGATGACCAGTACGGGGATGGGGAGAATAAAT | 0.41% | CGCAGCACGGGCGGCTCACGTACACCTGGAAGGAGATCGACGT | 0.40% |
| | mutation4 | ACAATCAACACCGATGACCAGTACGGGGATGGAGAGAATAAAC | 0.41% | CGCAGCACGAGCGGCTCACGTACGCCTGGAAGGAGATCGACGT | 0.39% |
| | mutation5 | ACAATCAACACCGATGACCAGTACGGGGATGGAGGGAATAAAT | 0.40% | CGCAGCACGAGCGGCTCACGTACACCTGGAGGGAGATCGACGT | 0.38% |
| | mutation6 | ACGATCAACACCGATGACCAGTACGGGGATGGAGAGAATAAAT | 0.40% | CGCAGCACGAGCGGCTCACGTACACCTGGAAGGGGATCGACGT | 0.36% |
| | mutation7 | GCAATCAACACCGATGACCAGTACGGGGATGGAGAGAATAAAT | 0.37% | CGCGGCACGAGCGGCTCACGTACACCTGGAAGGAGATCGACGT | 0.35% |
| | mutation8 | ACAATCAACACCGATGGCCAGTACGGGGATGGAGAGAATAAAT | 0.35% | CGCAGCACGAGCGGCTCACGTACACCTGGAAGGAGATCGGCGT | 0.33% |
| | mutation9 | ACAATCAACACCGATGACCAGTACGGGGGTGGAGAGAATAAAT | 0.35% | CGCAGCACGAGCGGCTCACGCACACCTGGAAGGAGATCGACGT | 0.33% |
| | mutation10 | ACAATCAACGCCGATGACCAGTACGGGGATGGAGAGAATAAAT | 0.35% | CGCAGCGCGAGCGGCTCACGTACACCTGGAAGGAGATCGACGT | 0.31% |

j. Edit events at $gRNA^{Aswhite}$ sites in $F_1$ ovaries (white eye $F_1$ from *Asvasa2-Cas9* line ♀ × *AsU6-2-gRNA*$^{Aswhite}$ ♂)

| | | gRNA$^{Aswhite1}$ | | gRNA$^{Aswhite2}$ | |
|---|---|---|---|---|---|
| | SeqName | Sequence features | Frequency | Sequence features | Frequency |
| | wild type | ACAATCAACACCGATGACCAGTACGGGGATGGAGAGAATAAAT | 6.16% | CGCAGCACGAGCGGCTCACGTACACCTGGAAGGAGATCGACGT | 24.62% |
|  | mutation1 | ACAATCAACACCGAT-----------GGATGGAGAGAATAAAT | 18.05% | CGCAGCACGAGCGGCTC----------GGAAGGAGATCGACGT | 13.69% |
|  | mutation2 | ACAATCAACACCGATG--------GGGGATGGAGAGAATAAAT | 9.73% | CGCAGC--------------------GGAAGGAGATCGACGT | 10.37% |
|  | mutation3 | ACAATCAACACCGATCGGATGATCCACCGGATGTGGGGATGGAGA GAATAAAT | 7.69% | CGCAGCACGAGCGGCTCACG------TGGAAGGAGATCGACGT | 5.03% |
|  | mutation4 | ACAATCAACACCGATG-------CGGGGATGGAGAGAATAAAT | 3.97% | CGCAGCACGAGCGGCTCAC--------GGAAGGAGATCGACGT | 4.22% |
| individual 1 | mutation5 | ACAATCAACACCGATG-CCAGTACGGGGATGGAGAGAATAAAT | 3.87% | CGCAGCACGAGCGGCTCACGTACACCTTGGAAGGAGATCGACGT | 2.92% |
|  | mutation6 | ACAATCAACACCGATG----GTACGGGGATGGAGAGAATAAAT | 3.63% | CGCAGCACGAGCGGC-----------GGAAGGAGATCGACGT | 2.83% |
|  | mutation7 | ACAATCAAC-------ACCAGTACGGGGATGGAGAGAATAAAT | 3.41% | CGCAGCACGAGCGG-----------TGGAAGGAGATCGACGT | 1.58% |
|  | mutation8 | ACAATCAACACCGATG--CAGTACGGGGATGGAGAGAATAAAT | 3.10% | CGCA---------------------TGGAAGGAGATCGACGT | 1.58% |
|  | mutation9 | ACAATCAACACCGAT-----GTACGGGGATGGAGAGAATAAAT | 2.89% | CGCAGCACGAGCGGCTCACGTACAC--GGAAGGAGATCGACGT | 1.56% |
|  | mutation10 | ACAATCAACACCGATG-------------------------T | 2.71% | CGCAGCACGAGCGGCTCACGTACACC-GGAAGGAGATCGACGT | 1.56% |
| | wild type | ACAATCAACACCGATGACCAGTACGGGGATGGAGAGAATAAAT | 4.64% | CGCAGCACGAGCGGCTCACGTACACCTGGAAGGAGATCGACGT | 15.48% |
|  | mutation1 | ACAATCAACACCGAT-----------GGATGGAGAGAATAAAT | 7.59% | CGCAGCACGAGCGGCTCAC--------GGAAGGAGATCGACGT | 11.57% |
|  | mutation2 | ACAATCAACACCGATG--------GGGGATGGAGAGAATAAAT | 7.22% | CGCAGCACGAGCGGCTCACG------TGGAAGGAGATCGACGT | 6.25% |
| | mutation3 | ACAATCAACACCGATG-CCAGTACGGGGATGGAGAGAATAAAT | 6.35% | CGCAGCACGAGCGGCTCACGTAAGTCACGTACACGAAGGAGATCG ACGT | 4.95% |
|  | mutation4 | ACAATCAACACCGATGGACCAGTACGGGGATGGAGAGAATAAAT | 5.82% | CGCAGCACGAGCGGC-----------GGAAGGAGATCGACGT | 4.71% |
| individual 2 | mutation5 | ACAATCAACACCGATGTCACATGAACACCGATGTCACCGATGAACA GTACGGGGATGGAGAGAATAAAT | 5.21% | CGCAGCACGAGCGGCTCACGTACACCTTGGAAGGAGATCGACGT | 4.06% |
|  | mutation6 | ACAATCAACACCGATGTACGGTCAGTACGGGGATGGAGAGAATAA AT | 5.20% | CGCAGCACGAGCGGCTC----------GGAAGGAGATCGACGT | 3.99% |
|  | mutation7 | ACAATCAACACCGAT---------GGGGATGGAGAGAATAAAT | 4.65% | CGCAGCACGAGCGG----------CTGGAAGGAGATCGACGT | 3.38% |
| | mutation8 | ACAATCAACACCGATGACCAGTACGGGGATGGAGAGAATAAAT | 4.64% | CGCAGCACGAGCGGCTCACGTACA-CTGGAAGGAGATCGACGT | 2.38% |
| | mutation9 | ACAATCAACACCGATGGGGGACCAGTACGGGGATGGAGAGAATAAAT | 3.47% | CGCAGCACGAGCGGCTCACGTACAC--GGAAGGAGATCGACGT | 2.16% |
|  | mutation10 | ACAATCAACACCGAT------------GATGGAGAGAATAAAT | 3.03% | CGCAGCACGAGCGGCTCACGTACACC-GGAAGGAGATCGACGT | 1.96% |

k. Edit events at *gRNA*$^{Aswhite}$ sites in $F_1$ ovaries (mosaic eye $F_1$ from *Asvasa2-Cas9* line♀ × *AsU6-2-gRNA*$^{Aswhite}$♂)

| | | gRNA$^{Aswhite1}$ | | gRNA$^{Aswhite2}$ | |
|---|---|---|---|---|---|
| | SeqName | Sequence features | Frequency | Sequence features | Frequency |
| | wild type | ACAATCAACACCGATGACCAGTACGGGGATGGAGAGAATAAAT | 16.12% | CGCAGCACGAGCGGCTCACGTACACCTGGAAGGAGATCGACGT | 35.05% |
|  | mutation1 | ACAATCAACAC------------CGGGGATGGAGAGAATAAAT | 9.13% | CGCAGCACGAGCGGCTCACGTATCCGTGGAAGGAGATCGACGT | 10.93% |
| | mutation2 | ACAATCAACACCGAT-ACCAGTACGGGGATGGAGAGAATAAAT | 8.01% | CGCAGCACGAGCGGCTCACGTACACCGCTGACCTGGGAAGGAGAT CGACGT | 9.77% |
|  | mutation3 | ACAATCAACACCGAT-----------GGATGGAGAGAATAAAT | 6.99% | CGCAGCACGAGCGGCTCACGTACAC--GGAAGGAGATCGACGT | 5.38% |
| | mutation4 | ACAATCAACACCGATCGATCAGTACGGGGATGGAGAGAATAAAT | 5.18% | CGCAGCACGAGCGGCTCACGT---CACGGAAGGAGATCGACGT | 4.24% |
| individual 1 | mutation5 | ACAATCAACACCGATGGTGTACGGGGAACGATGGTGTACGGAACC AGTACGGGGATGGAGAGAATAAAT | 4.28% | CGCAGCACGAGCGGCTC----------GGAAGGAGATCGACGT | 3.24% |
|  | mutation6 | ACAATCAACACCGATG--------GGGGATGGAGAGAATAAAT | 3.49% | CGCAGCACGAGCGGC-----------GGAAGGAGATCGACGT | 2.65% |
|  | mutation7 | ACAATCAACACCGATG-CCAGTACGGGGATGGAGAGAATAAAT | 2.83% | CGCAGCACGAGCGG------------GGAAGGAGATCGACGT | 1.90% |
|  | mutation8 | ACAATCAACACCGATGA-------GGGGATGGAGAGAATAAAT | 2.64% | CGCAGCACGAGCGGCTCACG------TGGAAGGAGATCGACGT | 1.83% |
|  | mutation9 | ACAATCAACACCGATG--CAGTACGGGGATGGAGAGAATAAAT | 2.33% | CGCAGCACGAGCGGCTCAC--------GGAAGGAGATCGACGT | 1.60% |
|  | mutation10 | ACAATCAACACCGATG------ACGGGGATGGAGAGAATAAAT | 1.99% | CGCAGCACGAGCGGCTCACGTACACCTTGGAAGGAGATCGACGT | 1.36% |
| | wild type | ACAATCAACACCGATGACCAGTACGGGGATGGAGAGAATAAAT | 4.67% | CGCAGCACGAGCGGCTCACGTACACCTGGAAGGAGATCGACGT | 20.94% |
|  | mutation1 | ACAATCAACACC----------------------GAATAAAT | 35.48% | CGCAGCACGAGCGGCTC----------GGAAGGAGATCGACGT | 8.62% |
|  | mutation2 | ACAATCAACACCGATGACCCCAGTACGGGGATGGAGAGAATAAAT | 10.75% | CGCAGCACGAGCGGCTCAC--------GGAAGGAGATCGACGT | 6.40% |
| | mutation3 | ACAATCAACACCGATGGTACCATCAGTACGGGGATGGAGAGAATA AAT | 9.54% | CGCAGCACGAGCGGCTCACGTACACCTTGGAAGGAGATCGACGT | 3.97% |
|  | mutation4 | ACAATCAAC-------ACCAGTACGGGGATGGAGAGAATAAAT | 5.43% | CGCAGCACGAGCGGCTCACG------TGGAAGGAGATCGACGT | 3.39% |
| individual 2 | mutation5 | ACAATCAACACCGATG--------GGGGATGGAGAGAATAAAT | 2.94% | CGCAGCACGAGCGG----------CTGGAAGGAGATCGACGT | 3.22% |
|  | mutation6 | ACAATCAACACCGATG-------CGGGGATGGAGAGAATAAAT | 2.04% | CGCAGCACGAGCGGCTCACGTACACCCGGGAACGGAAGGAGATC GACGT | 3.22% |
|  | mutation7 | ACAATCAACACCGAT-----------GGATGGAGAGAATAAAT | 1.42% | CGCAGCACGAGCGGCT---------CTGGAAGGAGATCGACGT | 2.89% |
|  | mutation8 | ACAATCAACACCGATG---AGTACGGGGATGGAGAGAATAAAT | 1.29% | CGCAGCACGAGCGGCTCA-------CTGGAAGGAGATCGACGT | 2.32% |
|  | mutation9 | ACAATCAACACCGAT----------GGGATGGAGAGAATAAAT | 1.00% | CGCAGCACGAGCGGCT----------TGGAAGGAGATCGACGT | 2.13% |
|  | mutation10 | ACAATCAACACCGAT---------GGGGATGGAGAGAATAAAT | 0.97% | CGCAGCACGAGCGG-----------TGGAAGGAGATCGACGT | 1.69% |

l. Edit events at *gRNA*$^{Aswhite}$ sites in $F_1$ ovaries (mosaic eye $F_1$ from *Asvasa2-Cas9* line♂ × *AsU6-2-gRNA*$^{Aswhite}$♀)

| | | gRNA$^{Aswhite1}$ | | gRNA$^{Aswhite2}$ | |
|---|---|---|---|---|---|
| | SeqName | Sequence features | Frequency | Sequence features | Frequency |
| | wild type | ACAATCAACACCGATGACCAGTACGGGGATGGAGAGAATAAAT | 31.88% | CGCAGCACGAGCGGCTCACGTACACCTGGAAGGAGATCGACGT | 60.48% |
|  | mutation1 | ACAATCAACACCGATG--------GGGGATGGAGAGAATAAAT | 8.54% | --------------------------GGAAGGAGATCGACGT | 3.31% |
| | mutation2 | ACAATCAACACCGATG-CCAGTACGGGGATGGAGAGAATAAAT | 7.10% | CGCAGCACGAGCGGCTCACGTACA-CTGGAAGGAGATCGACGT | 2.15% |
|  | mutation3 | ACAATCAACACCG-----------GGGGATGGAGAGAATAAAT | 7.05% | CGCAGCACGAGCGGCTCACGTACAC--GGAAGGAGATCGACGT | 1.72% |
| | mutation4 | ACAATCAACACCGATG--CAGTACGGGGATGGAGAGAATAAAT | 3.36% | CGCAGCACGAGCGGCTCACGTACACC-GGAAGGAGATCGACGT | 1.71% |
| individual 1 | mutation5 | ACAATCAACACCGATG----------------GAGAATAAAT | 3.31% | CGCAGCACGAGCGGCTCACGTACACCTTGGAAGGAGATCGACGT | 1.63% |
|  | mutation6 | ACAATCAAC-------ACCAGTACGGGGATGGAGAGAATAAAT | 1.78% | CGCAGCACGAGCGGCTCACG------TGGAAGGAGATCGACGT | 1.46% |
|  | mutation7 | ACAATCAACACCGATG---AGTACGGGGATGGAGAGAATAAAT | 1.55% | CGCAGCACGAGCGGCTC----------GGAAGGAGATCGACGT | 1.35% |
|  | mutation8 | ACAATCAACACCG-----------------------ATAAAT | 1.46% | CGCAGCACGAGCGGCTCACGTAC----GGAAGGAGATCGACGT | 1.21% |
|  | mutation9 | ACAATCAACACCGAT-----------GGATGGAGAGAATAAAT | 1.39% | CGCAGCACGAGCGGCT----------TGGAAGGAGATCGACGT | 1.10% |
|  | mutation10 | ACAATCAACACC---GACCAGTACGGGGATGGAGAGAATAAAT | 0.99% | CGCAGCACGAGCGGCT---------CTGGAAGGAGATCGACGT | 0.83% |
| | wild type | ACAATCAACACCGATGACCAGTACGGGGATGGAGAGAATAAAT | 2.71% | CGCAGCACGAGCGGCTCACGTACACCTGGAAGGAGATCGACGT | 19.30% |
|  | mutation1 | ACAATCAACACCGATG--------GGGGATGGAGAGAATAAAT | 11.67% | CGCAGCACGAGCGGCTC----------GGAAGGAGATCGACGT | 6.60% |
|  | mutation2 | ACAATCAAC-------ACCAGTACGGGGATGGAGAGAATAAAT | 7.81% | CGCAGCACGAGCGGCT---------CTGGAAGGAGATCGACGT | 6.21% |
| | mutation3 | ACGATCAACACC---GACCAGTACGGGGATGGAGAGAATAAAT | 7.77% | CGCAGCACGAGCGGCTCACGTACACCTTGGAAGGAGATCGACGT | 5.38% |
|  | mutation4 | ACAATCAACACCGAT----------GGGATGGAGAGAATAAAT | 6.42% | CGCAGCACGAGCGG----------CTGGAAGGAGATCGACGT | 4.38% |
| individual 2 | mutation5 | ACAATCAACACCGATG-CCAGTACGGGGATGGAGAGAATAAAT | 4.62% | CGCAGCACGAGCGGCTCAC--------GGAAGGAGATCGACGT | 3.74% |
| | mutation6 | ACAATCAACACCGATG--CAGTACGGGGATGGAGAGAATAAAT | 4.20% | CGCAGCACGAGCGGCTCACGTACACC-GGAAGGAGATCGACGT | 2.79% |
|  | mutation7 | ACAATCAACACCGAT------------GATGGAGAGAATAAAT | 4.12% | CGCAGCACGAGCGGCTCACGCAC----GGAAGGAGATCGACGT | 2.74% |
|  | mutation8 | ACAATCAACACCGAT-----GTACGGGGATGGAGAGAATAAAT | 3.56% | CGCAGCACGAGCGGCTCACGTACAC--GGAAGGAGATCGACGT | 2.65% |
|  | mutation9 | ACAATCAACACCGATG------ACGGGGATGGAGAGAATAAAT | 2.77% | CGCAGCACGAGCGGCT----------TGGAAGGAGATCGACGT | 2.27% |
|  | mutation10 | ACAATCAACACCGAT----------GGATGGAGAGAATAAAT | 2.76% | CGCAGCACGAGCGGCTCACGTAC----GGAAGGAGATCGACGT | 1.92% |
