## supplemental figures tables and files for "Engineered Promoter System Enables High-efficiency Transgenic CRISPR Editing in Malaria Transmitting Mosquito *Anopheles sinensis*": Supplemental tables 20251001 NEW.pdf

Table S1 Inverse PCR results of the three *Cas9* transgenic lines

| | Features of partial sequences from inverse PCR results | Information of insertion site |
|---|---|---|
| *Asvasa2-Cas9* line-1* | ......CTAGGGTTAAAGGACCGCATGAAACAACGA TGAGAATATGGTAAGTA...... | Chr01, 105253628 |
| *Asvasa2-Cas9* line-2 | ......CTAGGGTTAAAGAATATAGCGAAGTGGAGA AATCTACAGCTGCTTGA...... | Chr01, 4408651 |
| *Aszpg-Cas9* line-1* | ......CTAGGGTTAAGTGAACAGGCAATAAAATTG GTCGCGATATTGATTGCA...... | Chr01, 14507875 |
| *Aszpg-Cas9* line-2 | ......CTAGGGTTAATGTGCATTTGCGCTTTACTGT GTTGCACTTTGGATCCGT...... | Chr01, 20472872 |
| *Asnanos-Cas9* line-1* | ......CTAGGGTTAAATAATCATGTCATGGCTTTTG TGCAATTTTCGCCATGAA...... | Chr02, 36568681 |
| *Asnanos-Cas9* line-2 | ......CTAGGGTTAAACAACAGTGACCTTCTCATTG AATGACGAATTCTTGCA...... | Chr01, 12101403 |
| *AsU6-1-gRNA$^{Aswhite}$* line-1* | ......CTAGGGTTAAAATACCAAACTGCCCACCAG CGGACTGTCGGGTGGAG...... | Chr01, 48411251 |
| *AsU6-1-gRNA$^{Aswhite}$* line-2 | ......CTAGGGTTAAAAGTTGTAAAGCTCTTTGCAA TCCACTAATTTTCCCC...... | Chr02, 86553888 |
| *AsU6-2-gRNA$^{Aswhite}$* line-1* | ......CTAGGGTTAAAGAAGTTCAAACCATGAAGG AAAGCCGAATTGAAACG...... | Chr02, 91304928 |
| *AsU6-2-gRNA$^{Aswhite}$* line-2 | ......CTAGGGTTAAACGTACAAGGTCAACGATCG CCACGTTGCCAGGGTTT...... | Chr02, 77327580 |

Note: The blue bases represent the sequence on the piggyBac plasmid, the red bases indicate the TTAA insertion sites, and the black bases correspond to the genomic sequence. * denotes the line retained for subsequent crossing experiments

Table S2 Editing efficiency of the *Aswhite* gene in $G_0$ embryos from crosses between *Cas9* transgenic females and the lab-reared *Aswhite* mutant males following *gRNA*$^{Aswhite}$ injection

| | mosaic | | wild type | | efficiency |
|---|---|---|---|---|---|
| | female | male | female | male | |
| | 52 | 45 | 35 | 32 | |
| *Asvasa2-Cas9* line | 53 | 48 | 36 | 35 | 59.9% ± 1.7% |
| | 60 | 52 | 33 | 36 | |
| | 6 | 8 | 69 | 80 | |
| *Aszpg-Cas9* line | 5 | 6 | 70 | 75 | 8.4% ± 1.3% |
| | 9 | 9 | 87 | 81 | |
| | 0 | 0 | 142 | 130 | |
| *Asnanos-Cas9* line | 0 | 0 | 136 | 135 | 0 |
| | 0 | 0 | 122 | 134 | |
| | 30 | 27 | 54 | 61 | |
| WT* | 31 | 29 | 72 | 70 | 32.8% ± 2.9% |
| | 44 | 42 | 82 | 75 | |

Note: *denotes $G_0$ embryos from crosses between wild type females and the lab-reared *Aswhite* mutant males following Cas9-*gRNA*$^{Aswhite}$ RNP injection

Table S3 Germline editing efficiency of $F_1$ female from crosses between mutant $G_0$ males (from *gRNA*$^{Aswhite}$ injection into *Cas9* × *Aswhite* mutant embryos) and lab-reared *Aswhite* mutant females.

| | white | wild type | efficiency |
|---|---|---|---|
| | 43 | 97 | |
| *Asvasa2-Cas9* line | 46 | 78 | 35.9% ±4.7% |
| | 51 | 77 | |
| | 0 | 128 | |
| *Aszpg-Cas9* line | 0 | 135 | 0 |
| | 0 | 158 | |
| | 0 | 165 | |
| *Asnanos-Cas9* line* | 0 | 154 | 0 |
| | 0 | 157 | |
| | 48 | 118 | |
| WT | 40 | 113 | 27.1% ± 1.5% |
| | 43 | 120 | |

Note: * denotes $F_1$ individuals in the *Asnanos-Cas9* group were generated by crossing $G_0$ male individuals with a normal phenotype (which may carry cryptic germline mutations) to the lab-reared *Aswhite* mutant females.

Table S4 $G_0$ editing efficiency of the *Aswhite* gene in the *Asvasa2-Cas9* line following injection of four *AsU6-gRNA*$^{Aswhite}$ plasmids

| | injected number | hatched number | pupation count | mosaic | | wild type | | efficiency |
|---|---|---|---|---|---|---|---|---|
| | | | | female | male | female | male | |
| | 902 | 86 | 39 | 13 | 11 | 8 | 7 | |
| *AsU6-1* | 852 | 87 | 51 | 15 | 16 | 10 | 10 | 62.1%±1.6% |
| | 859 | 98 | 61 | 17 | 22 | 9 | 13 | |
| | 1123 | 130 | 65 | 9 | 8 | 25 | 23 | |
| *AsU6-2* | 1073 | 121 | 67 | 10 | 13 | 20 | 24 | 31.1%±4.3% |
| | 987 | 105 | 58 | 8 | 11 | 19 | 20 | |
| | 892 | 162 | 89 | 30 | 26 | 15 | 18 | |
| *AsU6-3* | 732 | 144 | 76 | 21 | 23 | 16 | 16 | 60.2%±2.5% |
| | 902 | 105 | 62 | 17 | 20 | 13 | 12 | |
| | 826 | 83 | 45 | 13 | 15 | 7 | 10 | |
| *AsU6-4* | 911 | 86 | 42 | 13 | 11 | 9 | 9 | 59.9%±2.6% |
| | 1025 | 99 | 58 | 17 | 18 | 10 | 13 | |

Table S5 Editing efficiency of *Aswhite* following *AsU6*-1-*gRNA*$^{Aswhite}$ plasmid injection into embryos from *Aszpg-Cas9* or *Asnanos-Cas9* females crossed with lab-reared *Aswhite* mutant males.

| | mosaic | wild type | efficiency |
|---|---|---|---|
| | 15 | 164 | |
| *Aszpg-Cas9* line | 19 | 152 | 10.3± 1.6% |
| | 17 | 134 | |
| | 0 | 146 | |
| *Asnanos-Cas9* line | 0 | 167 | 0 |
| | 0 | 141 | |

Table S6 Phenotypic statistics of $F_1$ offspring from crosses between the three *Cas9* lines and the *AsU6*-1-*gRNA*$^{Aswhite}$ line

| | *Cas9* ♀ × *AsU6-1-gRNA*$^{Aswhite}$ ♂ (direct cross) | | | *Cas9* ♂ × *AsU6-1-gRNA*$^{Aswhite}$ ♀ (reciprocal cross) | | |
|---|---|---|---|---|---|---|
| | white | mosaic | wild type | white | mosaic | wild type |
| | 167 | 0 | 0 | 92 | 74 | 0 |
| *Asvasa2-Cas9* | 151 | 0 | 0 | 99 | 67 | 0 |
| | 158 | 0 | 0 | 93 | 70 | 0 |
| | 0 | 58 | 78 | 0 | 52 | 111 |
| *Aszpg-Cas9* | 0 | 55 | 71 | 0 | 56 | 116 |
| | 0 | 57 | 80 | 0 | 57 | 123 |
| | 0 | 0 | 141 | 0 | 0 | 184 |
| *Asnanos-Cas9* | 0 | 0 | 155 | 0 | 0 | 175 |
| | 0 | 0 | 133 | 0 | 0 | 167 |

Table S7 Editing efficiency of the *Aswhite* gene at two *gRNA* target sites in $F_1$ ovaries (*Cas9* × *AsU6-1-gRNA*$^{Aswhite}$)

| | Wild type | | Edited | | Total count | | Editing efficiency* | |
|---|---|---|---|---|---|---|---|---|
| | *gRNA*$^{Aswhite1}$ | *gRNA*$^{Aswhite2}$ | *gRNA*$^{Aswhite1}$ | *gRNA*$^{Aswhite2}$ | *gRNA*$^{Aswhite1}$ | *gRNA*$^{Aswhite2}$ | *gRNA*$^{Aswhite1}$ | *gRNA*$^{Aswhite2}$ |
| $F_1$, white eye phenotype (from *Asvasa2-Cas9* ♀ × *AsU6-1-gRNA*$^{Aswhite}$ ♂) | 136 | 38 | 48555 | 40445 | 48691 | 40483 | 99.78% | 99.86% |
| | 65 | 75 | 38446 | 41636 | 38511 | 41711 | | |
| $F_1$, white eye phenotype (from *Asvasa2-Cas9* ♂× *AsU6-1-gRNA*$^{Aswhite}$ ♀) | 442 | 996 | 130773 | 99124 | 131215 | 100120 | 99.70% | 99.31% |
| | 89 | 159 | 44821 | 38701 | 44910 | 38860 | | |
| $F_1$, mosaic eye phenotype (from *Asvasa2-Cas9* ♂× *AsU6-1-gRNA*$^{Aswhite}$ ♀) | 176 | 605 | 113356 | 164620 | 113532 | 165225 | | |
| | 229 | 403 | 44673 | 40523 | 44902 | 40926 | | |
| $F_1$, mosaic eye phenotype (from *Aszpg-Cas9* ♀ × *AsU6-1-gRNA*$^{Aswhite}$ ♂) | 25421 | 36961 | 17125 | 10807 | 42546 | 47768 | 42.14% | 24.18% |
| | 26726 | 31333 | 22198 | 16025 | 48924 | 47358 | | |
| $F_1$, wild-type eye phenotype (from *Aszpg-Cas9* ♀ × *AsU6-1-gRNA*$^{Aswhite}$ ♂) | 66108 | 116171 | 36279 | 28982 | 102387 | 145153 | | |
| | 33620 | 50757 | 30455 | 12952 | 64075 | 63709 | | |
| $F_1$, mosaic eye phenotype (from *Aszpg-Cas9* ♂× *AsU6-1-gRNA*$^{Aswhite}$ ♀) | 97739 | 104372 | 55753 | 36919 | 153492 | 141291 | 39.59% | 25.42% |
| | 41821 | 41095 | 24274 | 11502 | 66095 | 52597 | | |
| $F_1$, wild-type eye phenotype (from *Aszpg-Cas9* ♂× *AsU6-1-gRNA*$^{Aswhite}$ ♀) | 71259 | 91951 | 47076 | 27360 | 118335 | 119311 | | |
| | 21813 | 39007 | 18253 | 17344 | 40066 | 56351 | | |
| $F_1$, wild-type eye phenotype (from *Asvnanos-Cas9* ♀ × *AsU6-1-gRNA*$^{Aswhite}$ ♂) | 114426 | 112832 | 18296 | 13993 | 132722 | 126825 | 14.56% | 11.33% |
| | 35522 | 33407 | 6444 | 4403 | 41966 | 37810 | | |
| $F_1$, wild-type eye phenotype (from *Asnanos-Cas9* ♂× *AsU6-1-gRNA*$^{Aswhite}$ ♀) | 96295 | 98184 | 14041 | 10657 | 110336 | 108841 | 13.67% | 12.92% |
| | 30594 | 36287 | 5240 | 5028 | 35834 | 31315 | | |

*Editing efficiency is defined as the mutation rate associated with the editing event. Mutation rates induced by editing events were corrected for the background error rate determined from wild-type amplicon sequencing.

Table S8. Offspring phenotypes from crosses of *Asvasa2-Cas9* × *AsU6*-1 $F_1$ females (direct: white-eyed; reciprocal: mosaic eye/white-eyed) with lab-reared *Aswhite* mutant males

| | F₂, white | | | F₂, mosaic | | | F₂, wild type | | |
|---|---|---|---|---|---|---|---|---|---|
| | direct | reciprocal | | direct | reciprocal | | Direct | reciprocal | |
| | (F₁, white) | (F₁, white) | (F₁, mosaic) | (F₁, white) | (F₁, white) | (F₁, mosaic) | (F₁, white) | (F₁, white) | (F₁, mosaic) |
| Dual fluorescence | 10 | 16 | 32 | 0 | 0 | 0 | 0 | 0 | 0 |
| | 15 | 21 | 39 | 0 | 0 | 0 | 0 | 0 | 0 |
| | 23 | 31 | 46 | 0 | 0 | 0 | 0 | 0 | 0 |
| Red fluorescence | 70 | 70 | 88 | 0 | 0 | 0 | 0 | 0 | 0 |
| | 82 | 80 | 95 | 0 | 0 | 0 | 0 | 0 | 0 |
| | 76 | 86 | 92 | 0 | 0 | 0 | 0 | 0 | 0 |
| Green fluorescence | 60 | 62 | 76 | 0 | 0 | 0 | 0 | 0 | 0 |
| | 77 | 89 | 79 | 0 | 0 | 0 | 0 | 0 | 0 |
| | 81 | 87 | 81 | 0 | 0 | 0 | 0 | 0 | 0 |
| None fluorescence | 8 | 17 | 53 | 0 | 0 | 0 | 0 | 0 | 0 |
| | 16 | 23 | 68 | 0 | 0 | 0 | 0 | 0 | 0 |
| | 20 | 27 | 59 | 0 | 0 | 0 | 0 | 0 | 0 |

Note: "direct" indicates crosses between *Asvasa2-Cas9* females×*AsU6*-1 males, while "reciprocal" indicates crosses between *Asvasa2-Cas9* males×*AsU6*-1 females.

TableS9. Offspring phenotypes from crosses of *Aszpg-Cas9* ×*AsU6*-1 $F_1$ females (direct: mosaic eye/wild type ; reciprocal: mosaic eye/wild type ) with lab-reared *Aswhite* mutant males

| | $F_2$,white | | | | $F_2$,mosaic | | | | $F_2$,wild type | | | |
|---|---|---|---|---|---|---|---|---|---|---|---|---|
| | direct | | reciprocal cross | | direct | | reciprocal | | direct | | reciprocal | |
| | ($F_1$, mosaic) | ($F_1$, WT) | ($F_1$, mosaic) | ($F_1$, WT) | ($F_1$, mosaic) | ($F_1$, WT) | ($F_1$, mosaic) | ($F_1$, WT) | ($F_1$, mosaic) | ($F_1$, WT) | ($F_1$, mosaic) | ($F_1$, WT) |
| Dual fluorescence | 81 | 68 | 41 | 55 | 9 | 0 | 0 | 2 | 40 | 52 | 66 | 59 |
| | 90 | 79 | 39 | 63 | 7 | 5 | 0 | 12 | 47 | 55 | 70 | 73 |
| | 79 | 81 | 45 | 73 | 8 | 4 | 0 | 10 | 63 | 59 | 86 | 69 |
| Red fluorescence | 49 | 76 | 46 | 62 | 0 | 0 | 0 | 0 | 60 | 66 | 63 | 82 |
| | 54 | 62 | 48 | 55 | 0 | 0 | 0 | 0 | 67 | 64 | 75 | 97 |
| | 58 | 60 | 49 | 57 | 0 | 0 | 0 | 0 | 69 | 71 | 68 | 91 |
| Green fluorescence | 70 | 52 | 42 | 70 | 0 | 0 | 0 | 0 | 86 | 83 | 68 | 84 |
| | 77 | 56 | 45 | 72 | 0 | 0 | 0 | 0 | 92 | 82 | 57 | 82 |
| | 84 | 66 | 51 | 67 | 0 | 0 | 0 | 0 | 84 | 68 | 78 | 101 |
| None fluorescence | 26 | 36 | 42 | 42 | 0 | 0 | 0 | 0 | 68 | 67 | 76 | 62 |
| | 31 | 40 | 50 | 45 | 0 | 0 | 0 | 0 | 80 | 74 | 72 | 44 |
| | 34 | 42 | 58 | 50 | 0 | 0 | 0 | 0 | 87 | 67 | 77 | 51 |

Note: “direct” indicates crosses between *Asvzpg-Cas9* females×*AsU6*-1 males, while “reciprocal” indicates crosses between *Aszpg-Cas9* males×*AsU6*-1 females.

Table S10. Offspring phenotypes from crosses of *Asnanos-Cas9* × *AsU6*-1 $F_1$ females (wild-type eye in both direct and reciprocal crosses) with lab-reared *Aswhite* mutant males

| | $F_{2,}$ white | | $F_{2,}$ mosaic | | $F_{2,}$ Wild type | |
|---|---|---|---|---|---|---|
| | direct ($F_1$, wild type) | Reciprocal ($F_1$, wild type) | direct ($F_1$, wild type) | Reciprocal ($F_1$, wild type) | direct ($F_1$, wild type) | Reciprocal ($F_1$, wild type) |
| Dual fluorescence | 2 | 0 | 1 | 0 | 103 | 63 |
| | 0 | 0 | 0 | 0 | 97 | 78 |
| | 0 | 0 | 0 | 0 | 105 | 77 |
| Red fluorescence | 0 | 0 | 0 | 0 | 112 | 79 |
| | 0 | 0 | 0 | 0 | 115 | 90 |
| | 0 | 0 | 0 | 0 | 103 | 84 |
| Green fluorescence | 0 | 0 | 0 | 0 | 63 | 78 |
| | 0 | 0 | 0 | 0 | 68 | 54 |
| | 0 | 0 | 0 | 0 | 64 | 49 |
| None fluorescence | 0 | 0 | 0 | 0 | 18 | 67 |
| | 0 | 0 | 0 | 0 | 20 | 49 |
| | 0 | 0 | 0 | 0 | 19 | 56 |

Note: "direct" indicates crosses between *Asnanos-Cas9* females×*AsU6*-1 males, while "reciprocal" indicates crosses between *Asnanos-Cas9* males×*AsU6*-1 females.

Table S11. Phenotype analysis of $F_1$ from *Asvasa2-Cas9* × *AsU6-2-gRNA*$^{Aswhite}$

| | white | mosaic | wild type | white eye proportion |
|---|---|---|---|---|
| *Asvasa2-Cas9* ♀ × *AsU6-2-gRNA*$^{Aswhite}$ ♂ | 50 | 82 | 0 | 36.8%±1.1% |
| | 51 | 87 | 0 | |
| | 47 | 85 | 0 | |
| *Asvasa2-Cas9* ♂ × *AsU6-2-gRNA*$^{Aswhite}$ ♀ | 0 | 121 | 0 | 0 |
| | 0 | 116 | 0 | |
| | 0 | 123 | 0 | |

TableS12. Editing efficiency of the *Aswhite* gene at two *gRNA* target sites in $F_1$ ovaries (*Asvasa2-Cas9* × *AsU6-2-gRNA*$^{Aswhite}$)

| | WT | | Edited | | Total-count | | Editing efficiency | |
|---|---|---|---|---|---|---|---|---|
| | $gRNA^{Aswhite1}$ | $gRNA^{Aswhite2}$ | $gRNA^{Aswhite1}$ | $gRNA^{Aswhite2}$ | $gRNA^{Aswhite1}$ | $gRNA^{Aswhite2}$ | $gRNA^{ASwhite1}$ | $gRNA^{Aswhite2}$ |
| $F_1$, white eye phenotype (from *Asvasa2-Cas9*♀ × *AsU6-2-gRNA*$^{Aswhite}$ ♂) | 6849 | 23415 | 104248 | 71705 | 111097 | 95120 | 94.60% | 79.95% |
| | 1932 | 5389 | 39741 | 29435 | 41673 | 34824 | | |
| $F_1$, mosaic eye phenotype (from *Asvasa2-Cas9*♀ × *AsU6-2-gRNA*$^{Aswhite}$ ♂) | 19987 | 35585 | 103977 | 65937 | 123964 | 101522 | 89.60% | 72.00% |
| | 1764 | 9994 | 36007 | 37734 | 37771 | 47728 | | |
| $F_1$, mosaic eye phenotype (from *Asvasa2-Cas9*♂× *AsU6-2-gRNA*$^{Aswhite}$♀) | 35223 | 68231 | 75275 | 44589 | 110498 | 112820 | 82.71% | 60.11% |
| | 1006 | 9202 | 36130 | 38483 | 37136 | 47685 | | |

*Editing efficiency is defined as the mutation rate associated with the editing event. Mutation rates induced by editing events were corrected for the background error rate determined from wild-type amplicon sequencing.

Table S13. Offspring phenotypes from crosses of *Asvasa2-Cas9* × *AsU6*-2 $F_1$ females (direct: white-eyed; reciprocal: mosaic/wild type) with lab-reared *Aswhite* mutant males

| | $F_2$, white | | | $F_2$, mosaic | | | $F_2$, wild type | | |
|---|---|---|---|---|---|---|---|---|---|
| | direct | | reciprocal | direct | | reciprocal | direct | | reciprocal |
| | ($F_1$, white) | ($F_1$, mosaic) | ($F_1$, mosaic) | ($F_1$, white) | ($F_1$, mosaic) | ($F_1$, mosaic) | ($F_1$, white) | ($F_1$, mosaic) | ($F_1$, mosaic) |
| Dual fluorescence | 26 | 55 | 82 | 14 | 21 | 37 | 0 | 0 | 0 |
| | 30 | 58 | 79 | 12 | 23 | 40 | 0 | 0 | 0 |
| | 32 | 56 | 77 | 14 | 31 | 45 | 0 | 0 | 0 |
| Red fluorescence | 46 | 80 | 74 | 8 | 16 | 76 | 0 | 0 | 2 |
| | 53 | 79 | 81 | 15 | 18 | 79 | 0 | 1 | 0 |
| | 55 | 81 | 82 | 12 | 13 | 65 | 0 | 0 | 0 |
| Green fluorescence | 40 | 60 | 46 | 30 | 61 | 14 | 0 | 1 | 3 |
| | 58 | 55 | 47 | 43 | 57 | 19 | 0 | 1 | 0 |
| | 51 | 62 | 53 | 42 | 64 | 23 | 0 | 0 | 0 |
| None fluorescence | 50 | 75 | 18 | 46 | 62 | 7 | 0 | 4 | 0 |
| | 60 | 68 | 19 | 54 | 50 | 11 | 0 | 2 | 1 |
| | 52 | 74 | 22 | 52 | 56 | 8 | 0 | 0 | 0 |

Note: "direct" indicates crosses between *Asvasa2-Cas9* females×*AsU6*-2 males, while "reciprocal" indicates crosses between *Asvasa2-Cas9* males×*AsU6*-2 females.

Table S14. Primers used in this study

| Primer names | Primer sequences (5'-3') | Purpose |
|---|---|---|
| *Asvasa2*-pro-F | AATGATCATCATGACAGATCTAGCCCGTTTGACATTTGCTG | Construction for *Asvasa2-Cas9* vector |
| *Asvasa2*-pro-R | CCATGGTGGCCTTTTAAATCGTGTGTCTCTGTAAC | |
| *Asvasa2-Cas9*-F | GATTTAAAAGGCCACCATGGACTATAAGGACC | |
| *Asvasa2-Cas9*-R | AGCGCACCATGCTACCGCTGCCGCTACC | |
| *Asvasa2-DsRed*-F | CAGCGGTAGCATGGTGCGCTCCTCCAAG | |
| *Asvasa2-DsRed*-R | TTCCACTCCACTACAGGAACAGGTGGTGGC | |
| *Asvasa2-ter*-F | GTTCCTGTAGTGGAGTGGAAGACGACTC | |
| *Asvasa2*-ter-R | GCACTGAACATTGTCAGATCTGACCCCAACGCATTTGTAG | |
| *Aszpg*-pro-F | AATGATCATCATGACAGATCTAACTGTTGTGAGGTATCTTCCTCT | Construction for *Aszpg-Cas9* vector |
| *Aszpg*-pro-R | CCATGGTGGCTTTGGGGCTAACCGCGCT | |
| *Aszpg*-Cas9-F | TAGCCCCAAAGCCACCATGGACTATAAGGACC | |
| *Aszpg*-DsRed-R | CCCTTACTGTCTACAGGAACAGGTGGTGGC | |
| *Aszpg*-ter-F | GTTCCTGTAGACAGTAAGGGATCTTAGAAAATGG | |
| *Aszpg*-ter-R | GCACTGAACATTGTCAGATCTAGAAGCAATTAACCAGCTCTCG | |
| *Asnanos*-pro-F | AATGATCATCATGACAGATCTGCAATACGCCCTTTCTGGTC | Construction for *Asnanos-Cas9* vector |
| *Asnanos*-pro-R | CCATGGTGGCGTTTTGATTTGATGTGGGTAAAATG | |
| *Asnanos*-Cas9-F | AAATCAAAACGCCACCATGGACTATAAGGACC | |
| *Asnanos*-DsRed-R | GCTCTTCTTTCTACAGGAACAGGTGGTGGC | |
| *Asnanos*-ter-F | GTTCCTGTAGAAAGAAGAGCAATTGTCTGGAAC | |
| *Asnanos*-ter-R | GCACTGAACATTGTCAGATCTCAACCTCCAGCAAGAATTAAAAG | |
| *AsU6*-1-F | AATGATCATCATGACAGATCTTGGAAATTACTTAATGCGCCGTGG | Construction for four *AsU6-gRNA$^{white}$* vectors |
| *AsU6*-1-R | ATGACCAGTACGGGGATGGACAAGCAGAGGCTGTTTGAAA | |
| *AsU6*-2-F | AATGATCATCATGACAGATCTACGCTGCTTCGATTGTGC | |
| *AsU6*-2-R | ATGACCAGTACGGGGATGGACGAAGCAAGGACATTTTGTT | |
| *AsU6*-3-F | AATGATCATCATGACAGATCTGTTCAATTTTTTCAACCCGCCT | |
| *AsU6*-3-R | ATGACCAGTACGGGGATGGACGAGGGACGTTCATTCGA | |
| *AsU6*-4-F | AATGATCATCATGACAGATCTAAATAAAAACAGCTAAATAAACATC | |
| *AsU6*-4-R | ATGACCAGTACGGGGATGGACGAGGGACGTTCATTCGA | |
| *gRNA-white*-scaffold-F | TCCATCCCCGTACTGGTCATGTTTTAGAGCTAGAAATAGC | |
| *gRNA-white*-scaffold-R | GCACTGAACATTGTCAGATCTAAAAAAAAGCACCGACTCGGTG | |
| *AsU6-1-gRNA*-F | TGTGGTCTCTCTTGTCCATCCCCGTACTGGTCATGTTTTAGAGCTAGAAATAGCA | Construction for *piggyBac-AsU6-gRNA$^{white}$* vectors (for engineering of *U6-gRNA$^{white}$* transgenic line) |
| *AsU6-1-gRNA*-R | TGTGGTCTCGAAACTCCAGGTGTACGTGAGCCGCCAAGCAGAGGCTGTTTGAAATTGT | |
| *AsU6-2-gRNA*-F | TGTGGTCTCTTTCGTCCATCCCCGTACTGGTCATGTTTTAGAGCTAGAAATAGCA | |

| | | |
|---|---|---|
| *AsU6-2-gRNA*-R | TGTGGTCTCGAAACTCCAGGTGTACGTGAGCCGCCGAAGCAA GGACATTTTGTTTCGT | |
| 6-mer | NNNNNN | Universal primer for 3' RACE |
| GeneRacer-OligodT | GCTGTCAACGATACGCTACGTAACGGCATGACAGTGTTTTTTT TTTTTTTTTTTTTTTT | |
| GeneRacer-3-R | GCTGTCAACGATACGCTACGTAACG | |
| GeneRacer-3-Nested-R | CGCTACGTAACGGCATGACAGTG | |
| *Asvasa2*-generacer-F | AGTGGCGGCTATGGTGGTGGTTC | 3' RACE for *Asvasa2* |
| *Asvasa2*-nest-F | AGGAGCCTGAAGAAGAGTGGTAGT | |
| *Aszpg*-3generacer-F | CTCGGGCTTCTGGTGAGCGTTTC | 3' RACE for *Aszpg* |
| *Aszpg*-3nest-F | AGAGGAGGATGAAGCGGATGTTTGA | |
| *Asnanos*-3generacer-F | CCCATAAAACCCATCATCACCCCG | 3' RACE for *Asnanos* |
| *Asnanos*-3nest-F | CACCTGACTTATCTGCGAGCAACAAC | |
| *Aswhite*-target-site-F | GAAATTAATACGACTCACTATAGTCCATCCCCGTACTGGTCAT GTTTTAGAGCTAGAAATAGCAAG | Synthesis of *Aswhite* sgRNA |
| sgRNA-r | AAAAGCACCGACTCGGTGCCACTTTTTCAAGTTGATAACGGAC TAGCCTTATTTTAACTTGCTATTTCTAGCTCTAAAAC | |
| *Aswhite*-detection-F | GTTTGCGAACGGGTCAGTC | Single mutation sites detection of the *Aswhite* |
| *Aswhite*-detection-R | TTCTCGGGAAACAGTAGCACC | |
| NGS-*Aswhite loci 1*-detection-F | CCTACACGACGCTCTTCCGATCTGTTTGCGAACGGGTCAGTC | High-throughput detection of *Aswhite* mutation sites |
| NGS-*Aswhite loci 1*-detection-R | GTTCCTTGGCACCCGAGAATTCCACTATCATAACGAGAACGCT TGTG | |
| NGS-*Aswhite loci 2*-detection-F | CCTACACGACGCTCTTCCGATCTACAACGCTGACGAACGATAA G | |
| NGS-*Aswhite loci 2*-detection-R | GTTCCTTGGCACCCGAGAATTCCATGCGAGGTAAAGCAGTGGC | |
| Inverse-PCR-F1 | GACGCATGATTATCTTTTACGTGAC | Inverse PCR primers |
| Inverse-PCR-R1 | TGACACTTACCGCATTGACA | |
| Inverse-nest PCR -F2 | GCGATGACGAGCTTGTTGGTG | |
| Inverse-nest PCR-R2 | TCCAAGCGGCGACTGAGATG | |
| qRT*Asvasa2*-F | AGACCAGGTTTCCGACCACTT | RT-qPCR for *Asvasa2* |
| qRT*Asvasa2*-R | CTCAACTCGCAATCAGCACATC | |
| qRT*Aszpg*-F | AACTGGGGTTTCCGCCTTTG | RT-qPCR for *Aszpg* |
| qRT*Aszpg*-R | AACATCCGCTTCATCCTCCTCTA | |
| qRT*Asnanos*-F | CCTTCGTGCGTCTGACAAATAA | RT-qPCR for *Asnanos* |
| qRT*Asnanos*-R | AAAACTCTGAACACTTCCTCCTGC | |
| qRT*Cas9*-F | ACGGCAGAAGGGAAACGAAC | RT-qPCR for *Cas9* |
| qRT*Cas9*-R | CAGCACATTGTCCAGATTAGCG | |
| qRT*AsRPL49*-F | GATGCTGCCCGTGTATCTGC | RT-qPCR for *AsRPL49* (Inner control) |
| qRT*AsRPL49*-R | TCAATCTGTCCGCTCATTTCGT | |
